# The hippocampus binds movements to their temporal position in a motor sequence

**DOI:** 10.1101/2022.12.20.521084

**Authors:** Nina Dolfen, Serena Reverberi, Hans Op de beeck, Bradley R. King, Genevieve Albouy

## Abstract

A plethora of daily motor tasks consist of sequences of movements or steps that need to be performed in a specific order. Yet, it remains unclear how the brain represents sequential motor actions in a way that preserves their temporal order. Here, we used multivoxel pattern similarity analysis of functional Magnetic Resonance Imaging (fMRI) data acquired during motor sequence practice to investigate whether the hippocampus, a brain region known to support temporal order in the non-motor memory domain, represents information about the temporal order of sequential motor actions. We also examined such representation in other regions of the motor network (i.e., the premotor cortex (PMC), supplementary motor area (SMA), anterior superior parietal lobule (aSPL) and striatum) known for their critical role in motor sequence learning. Our results show that hippocampal activation patterns carried information about movements in their learned temporal position in the sequence (i.e., movement-position binding), but not about movements or positions in random movement patterns. In contrast, other ROIs showed evidence of binding in the sequence as well as movement (M1, SMA, PMC, putamen and aSPL) and position (aSPL and PMC) coding in random movement patterns. Importantly, movement coding contributed to sequence learning patterns in M1, SMA and PMC but not in the putamen and aSPL, suggesting a specific involvement of these regions in movement-position binding. Altogether, our findings provide novel insight into the role of the hippocampus in the motor memory domain and point to its capacity to bind movements to their temporal position in a motor sequence. Our results also deepen our understanding of how striatal and cortical regions contribute to motor sequence learning via movement coding or movement-position binding.

**Significance Statement:** Consistent evidence collected over the last two decades indicates that the hippocampus - a brain structure traditionally associated to declarative memory - is critically involved in motor memory. Yet, the functional role and representational contribution of the hippocampus during motor learning remains to be elucidated. Using a multivariate functional MRI approach, we show here that the hippocampus binds movements to their temporal position in a learned sequence of actions. These results point towards the involvement of the hippocampus in preserving information about temporal order in motor memory - a process well described for declarative memories. We suggest that the ability of the hippocampus to encode temporal order during sequence learning is common across declarative and non-declarative memory systems.

## Introduction

Many of our everyday tasks, such as making a cup of tea or coffee, consist of repeated sequences of highly similar motor actions or steps. Learning and remembering the correct order of these motor actions is fundamental for successful task completion. Yet, it remains unclear how the brain encodes and represents sequential motor actions in a way that preserves their order. In the last decade, findings in the non-motor memory domain have indicated that the hippocampus plays a critical role in representing and preserving the order of sequential information (1). Neuroimaging studies revealed that multivoxel patterns of activation in the hippocampus are sensitive to the order of items and contain information about the temporal order of episodic events (2) as well as items in learned sequences (i.e., letters in (3) and objects in (4)). Such hippocampal patterns have also been found to predict subsequent memory of the temporal order of the learned items (2, 5). Importantly, it is currently unknown whether the hippocampus represents information about the order of actions in motor sequences.

Hippocampal recruitment during motor sequence learning has consistently been reported in the literature (6–18). Prior work investigating the functional role of the hippocampus in this process indicates that hippocampal activity during early learning supports the development of a spatial, effector-independent, representation of the motor sequence that encompasses the goal of the series of movements to be executed (6, 19). Recent research suggests that these spatial representations are reactivated during post-encoding offline episodes (16) and are linked to successful motor memory retention (6, 20). The potential role of the hippocampus in coding the temporal order of movements (i.e., binding movements to their temporal position in a sequence) has been suggested in a previous behavioral study (21) and there is magnetoencephalographic (MEG) evidence to suggest that the medial temporal lobe codes for an abstract template of ordinal position during motor sequence preparation (22). In the current study, we used multivoxel pattern similarity analysis of functional Magnetic Resonance Imaging (fMRI) data acquired during the execution of learned sequences to elucidate the functional role of the hippocampus in temporal order coding during motor sequence learning.

In brief, participants learned a fixed sequence of eight finger movements through repeated practice. Immediately following learning, participants were scanned while practicing the learned sequential pattern (i.e., motor sequence learning task) as well as random movement patterns (i.e., control task) using the same fingers but pressed in a random order. Given the specific role of the hippocampus in encoding and preserving order in other memory domains (1) and based on evidence that the hippocampus is critical for learning series of movements (19, 23), we predicted that hippocampal activation patterns would carry information about finger movements in their learned temporal position in the sequence (i.e., finger-position binding) above and beyond information about finger movements or positions alone. We also examined such coding in other regions of the motor network (i.e., the premotor cortex (PMC), supplementary motor area (SMA), anterior superior parietal lobule (aSPL) and striatum) that have recently been implicated in the encoding of sequence order during motor sequence learning (24–26).

## Results

As illustrated in Figure 1A, all participants completed a (1) learning, (2) fMRI and (3) retention session during which they performed a serial reaction time task (SRTT). Task instruction was to respond as fast as possible to the location of a cue that appeared on the screen (8 locations, 8 fingers, no thumbs; Fig. 1B). Depending on the task condition, the cues either followed a pseudorandom (RND SRTT, control task) or fixed, repeating sequential pattern (SEQ SRTT; motor sequence learning task). The repeating pattern in the SEQ condition consisted of an eight-element sequence of finger movements in which each finger was pressed once. In the RND condition, there was no repeating sequence, but each key was pressed once every eight elements.

**Figure 1.**
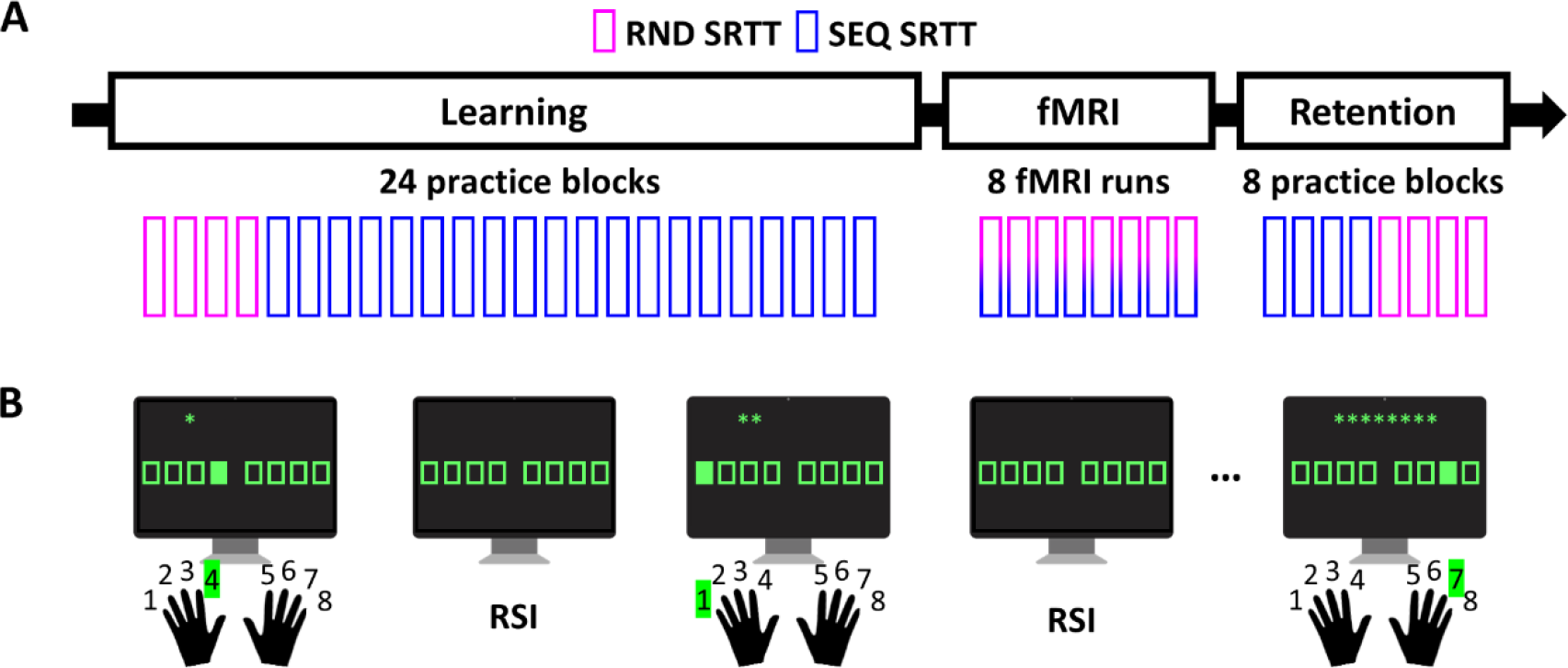
**(A) General procedure.** All participants completed a learning, fMRI and retention session during which they performed a serial reaction time task (SRTT; see panel B). Depending on the task condition, cues in the SRTT followed a random (RND) or repeating, sequential (SEQ) pattern. During the learning and retention sessions, the task took place outside the scanner and was organized in blocks of practice. During the fMRI session, data were acquired over eight consecutive scanning runs during which RND and SEQ SRTT conditions randomly alternated. **(B) Task.** During the SRTT, eight squares were presented on the screen, each corresponding to one of eight keys on the keyboard (no thumbs). Participants were instructed to press as quickly and accurately as possible the key corresponding to the location of a cue (green filled square; response to stimulus interval: 1.5-2.5s, average 2s). During both the RND and SEQ SRTT, the serial position of a trial in a stream of eight elements was indicated by green asterisks on top of the screen.

### Behavioral results

After baseline assessment of motor performance (see Supplementary Results section 2.1), participants performed the SEQ SRTT and learned a sequence of eight finger movements through repeated practice. Behavioral results showed a main effect of practice on performance speed (reaction time) and accuracy (% of correct responses) (Repeated Measures (RM) ANOVA; main effect of block; speed: F_(2.77,88.858)_ = 10.96, ɳ_p_^2^ = .255, *p* < .001; accuracy: F_(8.474,271.169)_ = 2.011, ɳ_p_^2^ = .059, *p* = .042; Fig. 2, learning). These results suggest that, as expected, participants learned the sequence of finger movements. The learning session was immediately followed by an fMRI session that captured the neural processes supporting the execution of the learned sequences. To optimize the multivariate analyses of the corresponding MRI data (see details in methods), RND and SEQ SRTT conditions were randomly alternated in each fMRI run. To investigate behavioral changes during the fMRI session, a 2 (condition) by 8 (run) RM ANOVA was performed on both speed and accuracy measures. Performance speed was significantly faster in the SEQ as compared to RND condition (condition: F(_1,31_) = 96.089, ɳ_p_^2^ = .756, *p* < .001) and fluctuated similarly across fMRI runs in both conditions (run: F_(2.518,78.061)_ = 3.246, ɳ_p_^2^ = .095, *p* = .034; run x condition: F_(4.882, 151.344)_ = .719, ɳ_p_^2^ = .023, *p* = .656; Fig. 2, fMRI). Performance accuracy was overall higher in the SEQ as compared to the RND condition (condition: F(_1,31_) = 26.908, ɳ_p_^2^ = .465, *p* < .001) and remained stable across runs in both conditions (run: F_(2.958,91.687)_ = .550, ɳ_p_^2^ = .017, *p* = .647; run x condition: F(_7, 217_) = .787, ɳ_p_^2^ = .025, *p* = .599; Fig. 2, fMRI). These results indicate that participants presented a sequence-specific performance advantage in the scanner despite the interleaved nature of practice. Last, results showed a main effect of condition on both speed and accuracy during retention, indicating sequence-specific retention of the skill outside of the scanning environment (condition x block RM ANOVA; condition: speed, F(_1,31_) = 199.082, ɳ_p_^2^ = .865, *p* < .001, accuracy, F(_1,31_) = 29.727, ɳ_p_^2^ = .490, *p* < .001; all other effects: all *p*s ≥ .172; Fig. 2, Retention).

**Figure 2.**
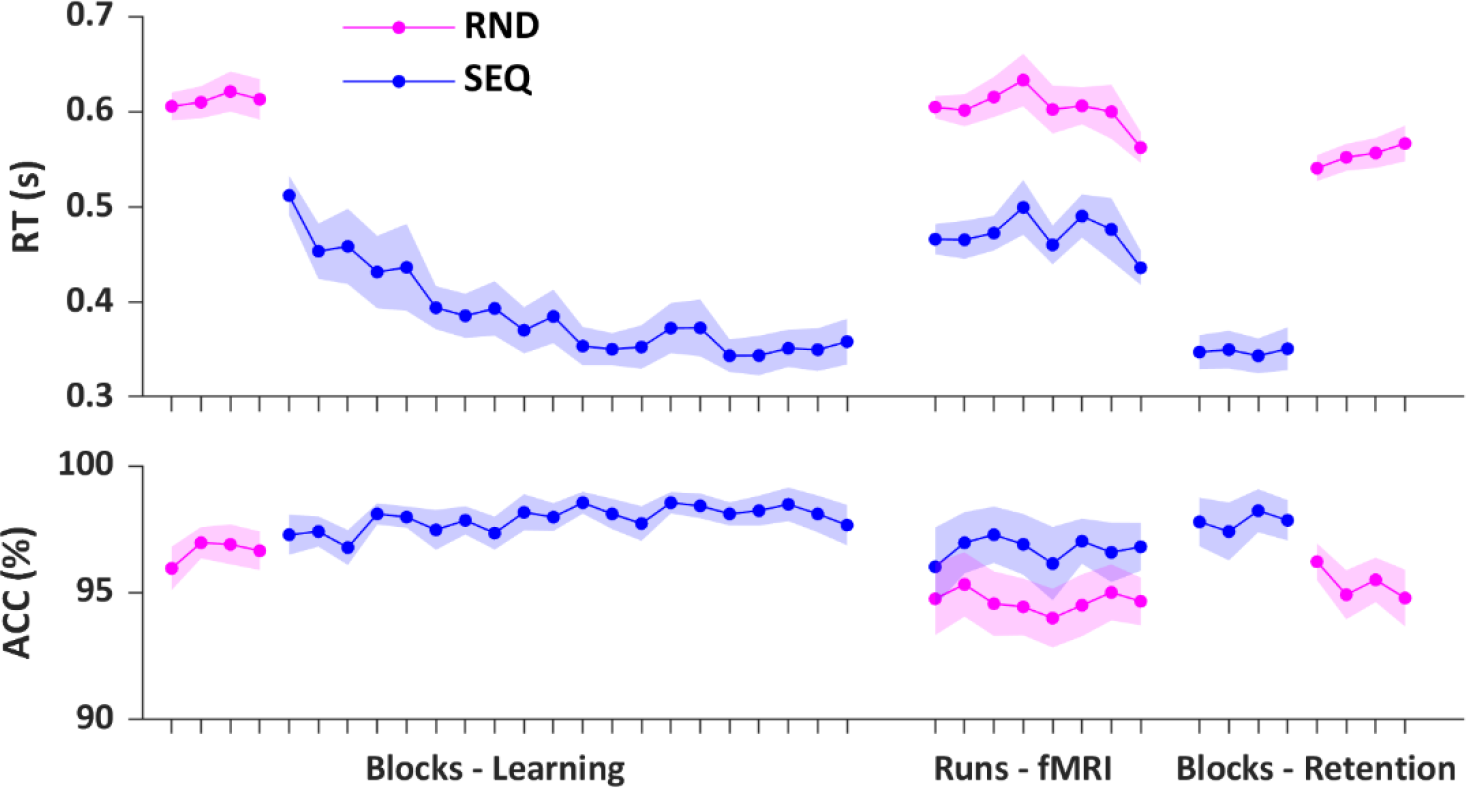
Behavioral results for all sessions. Reaction time (RT; s) and accuracy (ACC; %) for the RND and SEQ SRTT conditions are plotted across blocks of practice for the learning and retention sessions, and across runs for the fMRI session. N = 33. Shaded areas represent SEM

### Neuroimaging results

We used multivoxel pattern similarity analyses of task-related fMRI data to characterize coding of temporal order information during motor sequence execution. Similarity analyses are based on the notion that the relative pattern of activation across voxels in a given brain region is informative about the type of information that is processed by that region (27). Specifically, if a region codes for a particular type of information, higher similarity in voxel activity patterns is expected between pairs of trials sharing this information as compared to pairs of trials not sharing this information. Using this approach, we measured the extent to which specific regions of interest (ROIs) represent information about finger movements in their learned temporal position in the sequence (i.e., finger-position binding). Additionally, we performed control analyses using the random task to test whether the ROIs carried information about (1) finger movements irrespective of their position in the movement pattern (i.e., finger coding), and (2) temporal positions irrespective of the finger movement performed (i.e., position coding). Analyses were performed on seven bilateral ROIs involved in motor sequence learning: the primary motor cortex (M1), the supplementary motor cortex (SMA), the premotor cortex (PMC), the anterior part of the superior parietal lobule (aSPL), the hippocampus, the putamen, and the caudate nucleus (see regions of interest section in the methods).

#### Finger-position binding in a motor sequence

To investigate whether ROIs support the binding between finger movements and their learned temporal position in the sequence, we computed similarity in activation patterns between individual finger movements across repetitions of the motor sequence. This resulted in an 8×8 neural similarity matrix for each ROI (i.e., Sequence (SEQ) matrix; Fig. 3). Analysis of the ***SEQ matrix*** revealed significantly higher mean similarity along the diagonal (i.e., same finger + position) as compared to the off-diagonal (i.e., different finger + position) in all ROIs (paired sample t-test: diagonal SEQ vs. off-diagonal SEQ; hippocampus, t_32_ = 4.10, d = .71, *p*_FDR_ < .001; caudate, t_32_ = 3.88, d = .68, *p*_FDR_ < .001; putamen, t_32_ = 5.32, d = .96, *p*_FDR_ < .001; aSPL, t_32_= 10.01, d = 1.69, *p*_FDR_ <.001; SMA, t_32_ = 15.03, d = 2.62, *p*_FDR_ < .001; PMC, t_32_ = 17.30, d =3.05, *p*_FDR_ < .001; M1, t_32_ = 17.51, d = 3.05, *p*_FDR_ < .001; Fig. 4, SEQ matrix). These results suggest that activity patterns in all ROIs carry information about the binding between finger movements and their learned temporal position in the sequence. Importantly, as each finger movement in the sequence is bound to a specific position, both position and finger information are represented along the diagonal of the SEQ matrix. Accordingly, increased pattern similarity along the diagonal could be due to an overlap in either type of information. Control analyses were therefore performed using the random data to independently assess whether there is evidence of finger and/or position coding in our ROIs.

**Figure 3.**
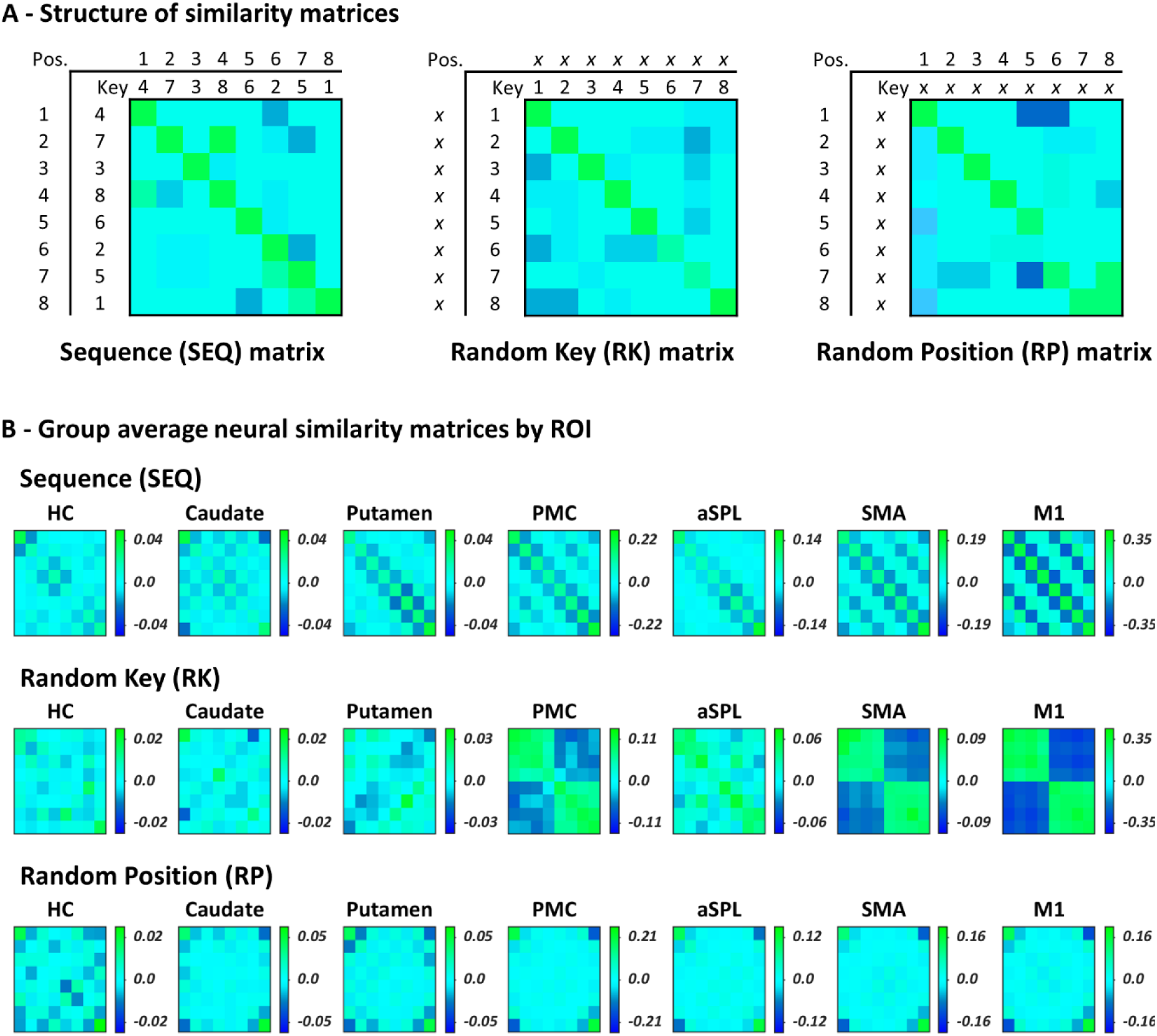
**(A)** Structure of neural similarity matrices for an exemplary sequence/individual. Pattern similarity was computed across repetitions of the motor sequence to assess finger-position binding in the sequence condition (SEQ matrix) as well as across repetitions of the random patterns to quantify coding of individual finger movements (i.e., key presses; Random Key (RK) matrix) and positions (Random Position (RP) matrix) in the random condition. Pairs of trials sharing a particular type of information (same finger + position, same finger, or same position) are represented on the diagonal of each matrix. **(B)** Group average neural similarity matrices for each ROI. Colour bars represent mean similarity (r). HC = Hippocampus. Note that colour scales are different between ROIs to accommodate for differences in signal-to-noise ratio (and therefore in effect sizes) between cortical and subcortical ROIs

**Figure 4.**
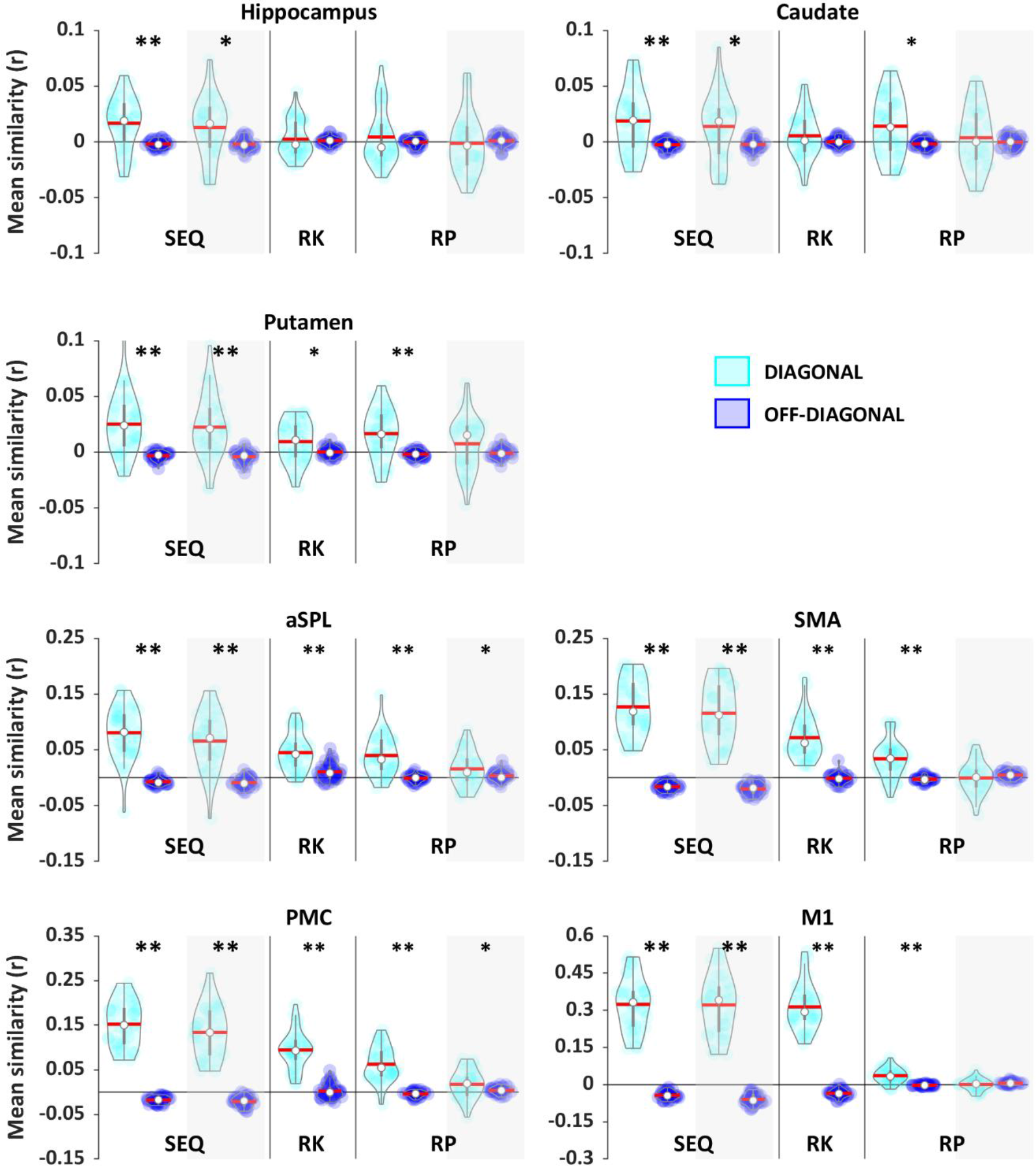
Mean pattern similarity for diagonal (cyan) and off-diagonal (blue) cells as a function of matrix and ROI. If a given ROI codes for a particular type of information, higher similarity is expected on the diagonal as compared to the off-diagonal. To assess whether ROIs code for movement-position binding in the sequence, we compared diagonal and off-diagonal cells in the Sequence (SEQ) matrix. To investigate whether ROIs carry information about movements (finger coding) and/or positions (position coding) in random movement patterns, diagonal and off-diagonal cells in the Random Key (RK) and in the Random Position (RP) matrices were compared. The hippocampus (HC) shows evidence for finger-position binding in the sequence condition but not for finger or position coding in the random condition. In contrast, other ROIs show evidence of finger-position binding in the sequence condition as well as position and/or finger coding during random practice. To verify whether results in the SEQ and RP matrices are driven by boundary effects, we reran the corresponding analyses without boundaries. Results after boundary correction are presented in the grey shaded areas. All ROIs show finger-position binding beyond boundaries in the sequence condition but only the PMC and aSPL carry information about positions (other than boundaries) under random conditions. Asterisks indicate significant differences between diagonal and off-diagonal (one sided paired sample t-test; \**pFDR*<.05 and ** *pFDR*<.001). Coloured circles represent individual data, jittered in arbitrary distances on the x-axis to increase perceptibility. Red horizontal lines represent means and white circles represent medians. Note that Y axis scales are different between ROIS to accommodate for differences in signal-to-noise ratio (and therefore in effect sizes) between cortical and subcortical ROIs.

#### Finger and position coding in random movement patterns

To assess finger coding irrespective of serial position, we quantified pattern similarity in the RND condition between repetitions of the same as well as different finger movements (i.e., key presses; Random Key (RK) matrix; Fig. 3). Consistent with a pattern of finger coding, analysis of the ***RK matrix*** showed that mean similarity on the diagonal (i.e., same finger) was significantly higher as compared to the off-diagonal (i.e., different finger) in M1, PMC, SMA, aSPL and putamen (paired sample t-test: diagonal RK vs. off-diagonal RK; M1, t_32_ = 18.93, d = 3.30, *p*_FDR_ < .001; PMC, t_32_ = 10.57, d = 1.84, *p*_FDR_ < .001; SMA, t_32_ = 9.85, d = 1.72, *p*_FDR_ < .001; aSPL, t_32_= 6.46, d = 1.12, *p*_FDR_ < .001; putamen, t_32_ = 2.46, d = .43, *p*_FDR_ = .01; Fig. 4). Exploratory follow-up analyses furthermore revealed higher similarity between different fingers belonging to the same as compared to different hands in these regions, which corresponds to a pattern consistent with hand coding (all t_32_ > 3.25, all d > .56, all *p*_FDR_ < .05; paired sample t-test: within-hand RK vs. between-hand RK; see methods for details). Note that within M1, aSPL, PMC and SMA, our results support the presence of finger coding beyond hand coding (M1, t_32_ = 9.944, d = 1.73, *p*_FDR_ < .001; PMC, t_32_ = 4.701, d = .82, *p*_FDR_ < .001; SMA, t_32_ = 2.693, d = .47, *p*_FDR_ = .01; aSPL, t_32_= 4.26, d = .74, *p*_FDR_ < .001; paired sample t-test: within-hand diagonal RK vs. within-hand off-diagonal RK). No evidence of finger coding was found in the hippocampus (t_32_ = .426, d = .07, *p*_FDR_ = .337) and caudate (t_32_ = 1.36, d = .24, *p*_FDR_ = .100).

To assess position coding, we computed pattern similarity between different finger movements sharing the same temporal position as well as between different finger movements in different temporal positions in the RND task condition (Random Position (RP) matrix; Fig. 3). Analysis of the ***RP matrix*** indicated that mean similarity was significantly higher for the diagonal (i.e., same position) as compared to the off-diagonal (i.e., different position) for all ROIs with the exception of the hippocampus (paired sample t-test: diagonal RP vs. off-diagonal RP; hippocampus, t_32_ = .940, d = .16, *p*_FDR_ = .185; caudate, t_32_ = 3.03, d = .52, *p*_FDR_ = .002; putamen, t_32_ = 4.33, d = .75*, p*_FDR_ < .001; aSPL, t_32_= 6.07, d= 1.06, *p*_FDR_ <.001; SMA, t_32_ = 5.56, d= .97, *p*_FDR_ < .001; PMC, t_32_ = 8.79, d = 1.53, *p*_FDR_ < .001; M1, t_32_ = 6.78, d = 1.18, *p*_FDR_ < .001; Fig. 4).

Altogether, these analyses indicate that hippocampal patterns carry information about finger movements in their learned temporal position in the sequence condition (i.e., finger-position binding), but not about finger movements or positions in the random condition. In contrast, other ROIs showed evidence of finger-position binding in the sequence condition as well as position (caudate, putamen, M1, SMA, PMC and aSPL; but see temporal position analyses below showing that higher similarity on positions 1 and 8 might account for these results in some ROIs) and finger/hand (putamen, M1, SMA, PMC and aSPL) coding during random practice.

#### Pattern similarity as a function of temporal position

Inspection of the SEQ and RND Position similarity matrices suggest that, in some regions, the increased pattern similarity along the diagonal highlighted in the previous analyses are not homogenous across serial positions but are driven by the boundaries of the 8-element pattern (i.e., higher similarity on positions 1 and 8; Fig. 3 and see Supplementary Results section 2.2 for corresponding statistical analyses). To verify whether the binding and position coding effects reported in the previous section are driven by these boundary effects, we excluded boundary positions from SEQ and RP matrices (see Fig. S1 for corresponding matrices) and reran the analyses. These control analyses showed that pattern similarity effects in the SEQ condition (same finger + position > different finger + position) remained significant in all ROIs (paired sample t-test: diagonal SEQ vs. off-diagonal SEQ; hippocampus, t_32_ = 2.66, d = .47, *p*_FDR_ = .013; caudate, t_32_ = 2.62, d = .46, *p*_FDR_ = .013; putamen, t_32_ = 4.47, d = .78, *p*_FDR_ < .001; aSPL, t_32_= 7.21, d = 1.26, *p*_FDR_ <.001; SMA, t_32_ = 12.5, d = 2.18, *p*_FDR_ < .001; PMC, t_32_ = 13.22, d = 2.30, *p*_FDR_ < .001; M1, t_32_ = 15.60, d = 2.71, *p*_FDR_ < .001; Fig. 4 – grey shaded areas), while there is no longer evidence that activation patterns in the caudate, putamen, M1 and SMA carry information about serial positions (other than boundary positions) in the random condition (paired sample t-test: diagonal RP vs. off-diagonal RP; caudate, t_32_ = .70, d = .12, *p*_FDR_ = .27; putamen, t_32_ = 1.67, d = .29, *p*_FDR_ = .073; M1, t_32_ = -1.18, d = -.21, *p*_FDR_ = .15; SMA, t_32_ = -.81, d = -.14, *p*_FDR_ = .25; Fig. 4 – grey shaded areas). The comparison in the random condition remained significant in PMC (t_32_ = 2.11, d = .37, *p*_FDR_ = .033) and aSPL (t_32_ = 1.9, d = .33, *p*_FDR_ = .049; Fig. 4 – grey shaded areas). In sum, these analyses confirm that all ROIs show finger-position binding beyond boundaries in the sequence condition. They also refine the classification described above and indicate that only the PMC and aSPL carry information about positions (other than boundaries) under random conditions. Altogether, these results indicate that hippocampal and caudate activation patterns carry information about finger-position binding but not about individual finger movements or positions in the random condition. In contrast, other ROIs showed evidence of binding in the sequence condition as well as finger (putamen, SMA, PMC and aSPL) and position (PMC and aSPL) coding during random practice. These conclusions are summarized in Table 1.

**Table 1.** Overview of results of the within-condition analyses

| Coding | Condition | Regions of interest |  |  |  |  |  |  |
| --- | --- | --- | --- | --- | --- | --- | --- | --- |
|  |  | HC | Caudate | Putamen | M1 | SMA | aSPL | PMC |
| Finger+position | SEQ | <.001 | <.001 | <.001 | <.001 | <.001 | <.001 | <.001 |
|  | w/o boundaries | .013 | .013 | <.001 | <.001 | <.001 | <.001 | <.001 |
| Finger | RND | .337 | .102 | .011 | <.001 | <.001 | <.001 | <.001 |
| Position | RND | .188 | .002 | <.001 | <.001 | <.001 | <.001 | <.001 |
|  | w/o boundaries | .342 | .270 | .073 | .150 | .250 | .049 | .033 |

**Notes.** Values are p-values based on paired sample t-tests (diagonal vs. off-diagonal). Boundaries refer to boundary positions 1 and 8 in an 8-element movement pattern. w/o = without.

#### Comparisons between representations

To assess whether ROIs showed preferential coding for one representation over the other, we compared delta similarity (i.e., mean pattern similarity diagonal minus off-diagonal) between conditions. For each ROI, delta values were entered in a RM ANOVA with condition (SEQ, RK and RP) as within-subjects factor. The analyses revealed a significant condition effect in all ROIs except the caudate (hippocampus, F(_2,64_) = 5.22, ɳ_p_^2^ = .14, *p_FDR_* = .009; caudate, F(_2,64_) = 2.73, ɳ_p_^2^ = .08, *p_FDR_* = .072; putamen, F(_2,64_) = 6.05, ɳ_p_^2^ = .16, *p_FDR_* = .005; M1, F(_2,64_) = 181.44, ɳ_p_^2^ = .85, *p_FDR_* < .001; PMC, F(_2,64_) = 52.87, ɳ_p_^2^ = .63, *p_FDR_* < .001; SMA, F(_2,64_) = 67.06, ɳ_p_^2^ = .68, *p_FDR_* < .001; aSPL, F(_2,64_) = 17.17, ɳ_p_^2^ = .35, *p_FDR_* < .001). Planned pairwise comparisons showed that delta similarity in the hippocampus was significantly larger in the SEQ as compared to the RK and RP conditions (SEQ vs. RK, *p_FDR_* = .005; SEQ vs. RP, *p_FDR_* = .05), while no difference was found between RP and RK (*p_FDR_* = .56) (Fig. 5A). These data suggest that the hippocampus supports finger-position binding beyond finger or position coding. A similar pattern of results was observed in aSPL (SEQ vs. RK, *p_FDR_* < .001; SEQ vs. RP, *p_FDR_* < .001; RK vs. RP, *p_FDR_* = .48; Fig. 5A). In the putamen finger-position binding was greater than finger coding (SEQ vs. RK, *p_FDR_* = .013) but no differences were found between the other conditions (SEQ vs. RP, *p_FDR_* = .08; RK vs. RP, *p_FDR_* = .08; Fig. 5A). Planned pairwise comparisons in M1 indicated no difference in delta similarity between SEQ and RK matrices (*p_FDR_* = .192), but both SEQ and RK delta similarity were significantly greater than RP (both *p_FDR_* <.001; Fig. 5A). Lastly, in PMC and SMA a gradient in delta similarity was observed such that SEQ was greater than RK which was greater than RP (SEQ vs. RK, *p_FDR_* < .001; SEQ vs. RP, *p_FDR_* < .001; RK vs. RP, *p_FDR_* < .001; Fig. 5A). In the caudate, results showed no difference in delta similarity between conditions (all *p_FDR_* > .1; Fig. 5A)

**Figure 5.**
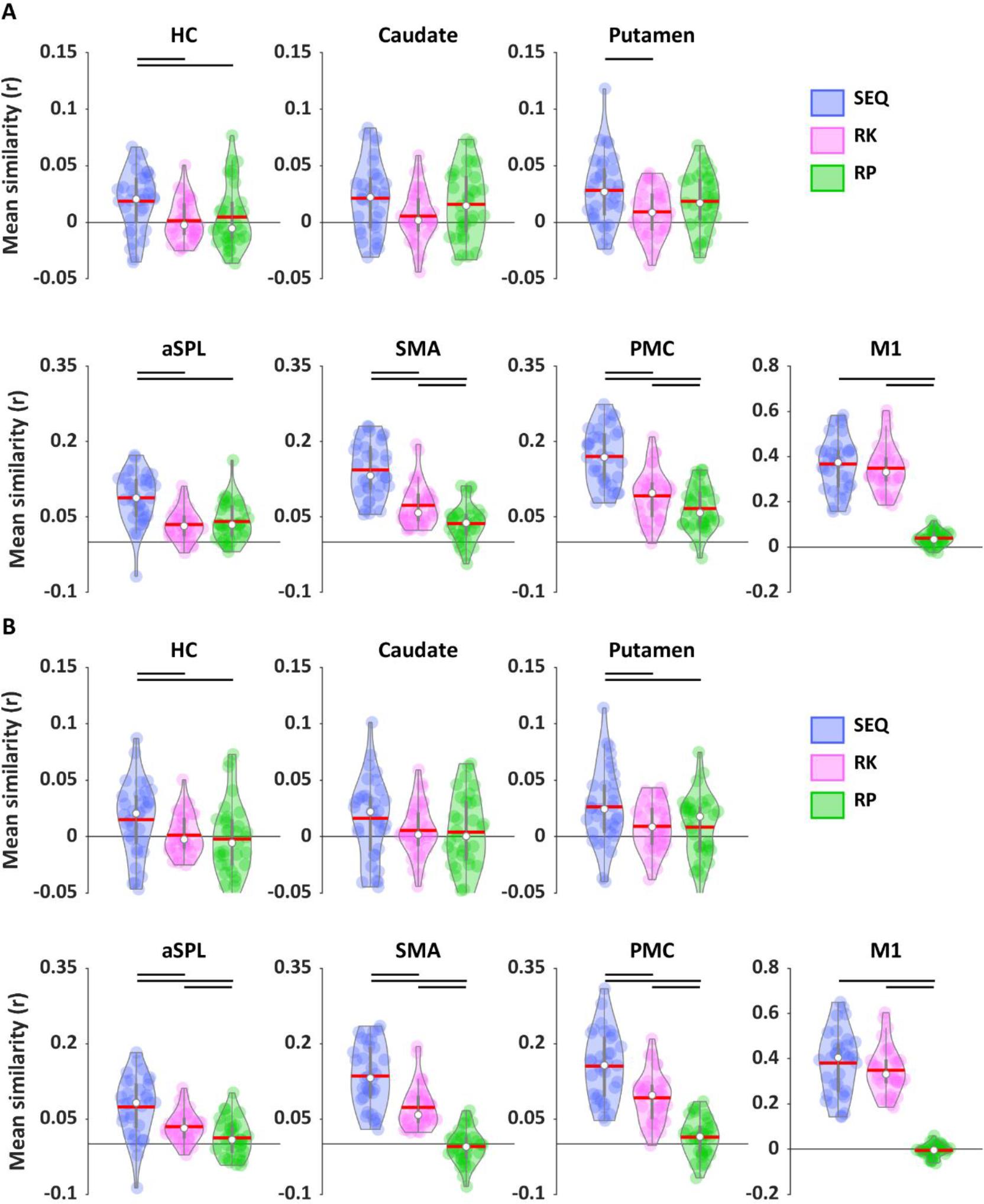
Delta pattern similarity (diagonal minus off-diagonal) as a function of condition and ROI. **(A)** The hippocampus (HC) supports finger-position binding beyond finger (SEQ > RK) and position coding (SEQ > RP), while the caudate shows no differences between conditions. The putamen shows binding beyond finger coding (SEQ > RK) and aSPL shows binding beyond finger (SEQ > RK) as well as position coding (SEQ > RP). In SMA and PMC binding is greater than finger coding (SEQ > RK) which was greater than position coding (RK > RP). M1 shows coding for binding and finger coding beyond position coding (SEQ > RP and RK > RP). **(B)** Results after boundary correction indicate that the putamen supports binding beyond finger and position coding, while aSPL supports finger beyond position coding. Horizontal lines indicate significant differences between conditions (planned pairwise comparisons following RM ANOVA; *pFDR* < .05). Coloured circles represent individual data, jittered in arbitrary distances on the x-axis to increase perceptibility. Red horizontal lines represent means and white circles represent medians. Note that Y axis scales are different between ROIS to accommodate for differences in signal-to-noise ratio (and effect sizes) between cortical and subcortical ROIs

For completeness, we reran these analyses while excluding the boundary positions from the SEQ and RP matrices. The control analyses showed a significant condition effect in all ROIs except the caudate (hippocampus, F(_2,64_) = 4.04, ɳ_p_^2^ = .11, *p_FDR_* = .025; caudate, F(_2,64_) = 1.53, ɳ_p_^2^ = .05, *p_FDR_* = .224; putamen, F(_2,64_) = 4.99, ɳ_p_^2^ = .14, *p_FDR_* = .014; M1, F(_2,64_) = 175.37, ɳ_p_^2^ = .85, *p_FDR_* < .001; PMC, F(_2,64_) = 75.45, ɳ_p_^2^ = .70, *p_FDR_* < .001; SMA, F(_2,64_) = 91.77, ɳ_p_^2^ = .74, *p_FDR_* < .001; aSPL, F(_2,64_) = 15.27, ɳ_p_^2^ = .32, *p_FDR_* < .001). Planned pairwise comparisons revealed the same pattern of results in all ROIs except for the putamen and aSPL (Fig. 5B). Specifically, when controlling for boundary effects, the putamen supports finger-position binding beyond finger (SEQ vs. RK, *p_FDR_* = .027) as well as position coding (SEQ vs. RP, *p_FDR_* = .027). The control analyses in aSPL revealed a similar gradient as in PMC and SMA such that SEQ was greater than RK which was greater than RP (SEQ vs. RK, *p_FDR_* = .002; SEQ vs. RP, *p_FDR_* = .012; RK vs. RP, *p_FDR_* < .001). The pattern in the other ROIs remained unchanged (hippocampus, SEQ vs. RK, *p_FDR_* = .038; SEQ vs. RP, *p_FDR_* = .038; RK vs. RP, *p_FDR_* = .56; caudate, all *p_FDR_* > .1; M1, SEQ vs. RK, *p_FDR_* = .192; SEQ vs. RP, *p_FDR_* < .001; RK vs. RP, *p_FDR_* < .001; PMC and SMA, SEQ vs. RK, *p_FDR_* < .001; SEQ vs. RP, *p_FDR_* < .001; RK vs. RP, *p_FDR_* < .001).

Altogether, these results indicate that the hippocampus and putamen show preferential coding for finger-position binding over position and finger coding. Interestingly, M1 showed preferential coding for both binding and finger (i.e., no difference between these two conditions) over position coding, while a gradient was observed in all the other cortical regions such that binding was greater than finger coding which was greater than position coding. The caudate nucleus did not show evidence of differential coding between conditions (i.e., no significant differences between conditions were found).

#### Contribution of finger and position coding to finger-position binding

Given that all ROIs presented evidence of finger-position binding and that some ROIs also coded for positions and/or finger movements, it seems possible that overlap in finger and/or position information might contribute to the enhanced pattern similarity for same finger+position in the learned sequence. Accordingly, in the last step, we used linear regression analyses to investigate whether finger and position coding (as assessed during random practice) might explain pattern similarity effects observed in the sequence condition. As expected in the hippocampus - where we observed no evidence of position or finger coding - neither factor contributed to the SEQ results (see Supplementary Table S3 for corresponding statistics). In the putamen and aSPL there was evidence for finger and/or position coding in the random task, however as for the hippocampus these factors did not contribute to the finger-position binding observed in the sequence task (see Table S3). Importantly, in M1, PMC and SMA, the analyses revealed that finger coding significantly contributed to the variance in SEQ delta similarity (M1, B = .565, t(_1,31_) = 2.337, *p_FDR_* = .021; PMC, B = .647, t(_1,31_) = 3.889, *p_FDR_* = .003; SMA, B = .540, t(_1,31_) = 2.573, *p_FDR_* = .035; see Table S3 for corresponding model statistics). Adding position to the model did not improve the model fit (see also Table S3 for linear regression models including only one independent variable showing that position coding does not contribute to sequence coding in these ROIs). Last, the finger-position binding observed in the caudate (that was not significantly greater than coding of other conditions) was not explained by either finger or position coding (Table S3).

In summary, the results of the linear regression models suggest that the involvement of the hippocampus, putamen and aSPL in the binding of movements to their temporal position in a sequence cannot be explained by position or movement coding. They also indicate that in M1, PMC and SMA finger-position binding during the execution of learned sequence can, at least in part, be explained by finger coding.

## Discussion

In the current study, we used multivoxel pattern similarity analysis of fMRI data to unravel the brain processes supporting the coding of temporal order information during the execution of learned motor sequences. In line with our hypothesis, we found that hippocampal multivoxel activation patterns specifically carried information about finger movements in their learned temporal position in the sequence (i.e., finger-position binding), but not about movements or positions in random movement patterns. In contrast, other ROIs showed evidence of binding in the sequence as well as position and/or movement coding in the random condition. Specifically, multivoxel patterns in M1, SMA and putamen carried movement-based information, while the PMC and aSPL represented both movement- and position-based information during random practice. Importantly, pattern similarity results in the sequence condition could, at least in part, be explained by finger coding in M1, PMC and SMA but not in the putamen and aSPL, suggesting a specific involvement of these regions in in movement-position binding in the sequence.

While hippocampal recruitment during motor sequence learning has consistently been reported in the literature (6–18), studies investigating the functional role of - and representational content in - the hippocampus during this process are sparse (but see (6, 16) for earlier research focusing on spatial representations of motor sequences in the hippocampus). Our analyses demonstrated that the hippocampus coded for finger movements in their learned temporal position in the sequence. As opposed to other ROIs, the hippocampus showed no evidence for movement or position coding and hence did not carry other task representations. These data suggest that the hippocampus specifically supports the binding between movements and their ordinal position in the sequence and thus the encoding of temporal order in the learned sequence. Our results are in line with a recent fMRI study showing that hippocampal multivoxel patterns differentiate between motor sequences based on the order of movements (17). They also concur with MEG evidence suggesting that the medial temporal lobe (the parahippocampus in particular) is involved in the coding of an abstract template of ordinal position during motor sequence preparation (22). More strikingly, the present findings concur with studies in the non-motor domain demonstrating that the hippocampus encodes the temporal order of objects (4) and auditory stimuli (3) during sequence learning. It is therefore tempting to speculate that the hippocampus processes temporal order information during sequence learning irrespective of the nature of the items (e.g., objects, sounds, movements) and the type of memory (i.e., motor, or non-motor).

In addition to the hippocampus, we also showed that finger-position binding in aSPL could not be explained by finger and/or position coding (even though this region showed evidence of such coding during random practice). Our data corroborate prior evidence that parietal regions encode the sequential context during motor learning (24–26) and hence are sensitive to the temporal order of movements. Together with the hippocampus, parietal areas are thought to support the spatial component of motor sequence learning acquired quickly during initial practice (19, 28). Based on earlier work in the motor domain showing that hippocampal and parietal regions support goal-based or spatial representations of motor sequences that are effector-independent (6), we hypothesize that binding processes in these regions might support the encoding of the temporal order of response goals or spatial targets in the learned sequence. Interestingly, such an associative map linking response goals to their ordinal position in the sequence was proposed to serve as a cognitive framework that can be abstracted during motor sequence learning and subsequently used as a cognitive-motor schema in which new movements can be rapidly integrated (21). In summary, our findings add to the growing body of literature associating hippocampo-cortical function to motor sequence learning (e.g., (6, 9, 15–17)) and indicate that the hippocampus and the parietal cortex represent temporal order information about motor sequences. They also suggest that the ability of the hippocampus to encode temporal order during sequence learning might be a general process that is shared across memory systems. This explanation, however, remains to be specifically tested.

A pattern of results corresponding to movement-position binding was also observed in the putamen. These findings are consistent with recent work showing that the execution of different motor sequences consisting of the same finger movements, but in a different order elicited distinct activation patterns in the striatum (17, 24). While these results appear to be similar in hippocampo-parietal areas and the striatum, we suggest that they support different processes. Specifically, the ability of the striatum to bind movements to their temporal position in a sequence might be related to its well-described role in the formation of stimulus-response associations (28–30). The striatum is indeed known to learn probabilistic or predictive associations between stimuli and responses, and/or between motor responses in a sequence (29, 31). The current data suggest that the striatum encoded the association between specific movements and their position in the learned deterministic sequence. Interestingly, our data also show that the putamen carried effector-specific information during random task practice. While these representations did not contribute to the binding process *per se*, previous literature has associated striatal responses to effector-dependent, motoric representation of the sequence during learning (28, 30, 32). We therefore argue that the striatal binding process might be movement-specific rather than goal-based as proposed for the hippocampus and parietal cortex. Given the rigid nature of stimulus-response associations learned by the putamen (31), we also speculate that the finger-position map developed in the striatum is less susceptible to be abstracted as compared to the hippocampo-cortical binding map described above. However, these statements are speculative and warrant further investigation.

Our data further indicate that the contribution of cortical regions such as the primary motor cortex (M1), premotor (PMC) and supplementary motor (SMA) to motor sequence learning might be attributed - at least in part - to movement coding. The results observed in M1 are consistent with earlier multivariate fMRI studies showing that M1 activation patterns relate to the execution of individual movements during motor sequence learning (25, 26). Importantly, our data demonstrate that movement-based information is not uniquely represented in the primary motor cortex during motor sequence learning but also in secondary motor areas (PMC and SMA). This is in contradiction with earlier work showing that the premotor and supplementary motor regions encode the sequential context rather than individual movement components (24–26) (but see (25) where no clear evidence for sequence representations was found in SMA). However, it is worth noting that while our analyses suggest a contribution via movement coding, they do not allow us to conclude that these regions do *not* code for finger-position binding during motor sequence learning (and hence do not represent sequence order). It is possible that regions code for multiple representations (e.g., both sequence order and individual movements; (33)). Yet, we argue that the predominant movement coding observed in the cortical ROIs in our research - in contrast to earlier work - might be attributed to differences in learning stages between studies. Whereas participants were extensively trained (up to 5 days) before the scanning session in previous work (24–26), participants only completed one hour of training in the current study. Taken together with evidence that sequence-specific activation patterns in secondary areas show less stability early in learning (24) and that the ability to discriminate between new sequences is significantly lower than between consolidated sequences in these areas (17), it could be argued that secondary motor areas support individual movement representations early during learning while sequence representations - and therefore movement-position binding - might emerge at later learning stages.

In addition to the movement coding discussed above, the premotor and parietal (aSPL) cortex showed a profile consistent with position coding during random task practice. While this effect was unexpected in the aSPL, the role of the premotor cortex in temporal processing of motor information is in line with our observations. Based on previous literature suggesting that the premotor cortex plays a key role in motor planning and action selection (e.g., (34–36)), it is tempting to speculate that position-specific multivoxel activation patterns in the PMC reflect action planning, i.e., the plan to complete a series of eight movements. However, as our analyses did not focus on this preparatory period, this interpretation needs to be made with caution. Nevertheless, we argue that PMC position patterns might reflect the availability of a temporal scaffold (here of eight elements) within which movements will be implemented. Such an interpretation is in line with prior research showing that the premotor cortex encodes temporal features about motor sequences (e.g., interval between key presses) irrespective of the movements performed (33). It is also in line with animal work indicating that PMC neurons are not only tuned to interval duration but also to specific positions in a series of movements (37). Although the implementation of such a position framework seems especially important to efficiently prepare and implement responses in the sequence task, these position coding processes were observed during random practice and did not significantly contribute to the binding pattern during sequence execution. It is possible that the visual cues indicating the ordinal position of movements in the 8-element stream (see Fig. 1B) might have triggered the development of such a framework in the random condition.

Given prior work suggesting that the SRTT is not exclusively a motor learning task but also includes important (non-motor) perceptual and explicit components (38, 39), it could be argued that the involvement of the hippocampus in the task is related to its perceptual and/or explicit components and therefore may not be generalizable to other motor sequence tasks without these attributes. We argue that this is unlikely for two reasons. First, significant hippocampal activations have been demonstrated during motor sequence learning independent of whether the task is visually guided or not (i.e., see (6, 7, 40–43)) for self-paced tasks without visual cues and (8, 10, 18, 44) for tasks with visual cues). Second, prior observations indicate that the hippocampus participates in the processing of sequential information in the motor domain irrespective of the awareness of the material to learn ((18) and see (19) for a review). Instead, we suggest that our findings might extend to other motor tasks in which sequential order of motor actions needs to be preserved. The contribution of the hippocampus across motor sequence learning tasks of different nature remains to be tested in future research.

Lastly, in all our ROIs, we found that pattern similarity was increased for boundary positions (i.e., positions 1 and 8) as compared to positions in the middle of the movement pattern in both task conditions. It is possible that the perception of boundaries was facilitated by the visual cues indicating the ordinal position of movements in the 8-element stream (i.e., asterisks on the top of the screen indicating the progress within the 8-element movement pattern; Fig. 1B). As these effects were observed in both the sequence and random tasks, we speculate that they might, at least in part, be driven by general attentional processes. Consistent with this idea, evidence in the non-motor domain shows that attentional states can modulate pattern similarity in cortical and subcortical regions (e.g., (45)). We suggest that attention was increased at the start and the end of the 8-element stream (i.e., either to adapt to/recognize the new condition or to prepare for a possible condition switch), which in turn modulated pattern similarity at these boundary positions. Finally, it seems possible that the lack of position and/or movement coding in the hippocampus is due to a generally lower signal-to-noise ratio in subcortical regions. However, given that there was evidence for effector and position (boundary) coding in striatal regions, this seems unlikely.

In summary, the present results demonstrate that the hippocampus, the parietal cortex and the striatum represent information about movements in their learned position in the sequence and hence encode the temporal order of movements. We propose that these regions might bind different representations, i.e., response goals in hippocampo-parietal areas as opposed to motor responses in the striatum, to their ordinal position in the sequence. Our data also refine the functional role of motor cortical regions (M1, SMA, PMC) in motor sequence learning and indicate a contribution through movement coding. Altogether, our results do not only provide novel insight into the role of the hippocampus in the procedural memory domain but also deepen our understanding of the involvement of striatal and cortical regions in motor sequence learning.

## Material and methods

### Participants

The study was approved by the Medical Ethics Committee of the University Hospital Leuven, Belgium (B322201525025). Thirty-five healthy, young (mean age: 23.4, range: 19-29, 20 females), right-handed (Edinburgh Handedness questionnaire; (46)) adults were recruited to participate in the study. All participants provided written informed consent at the start of the study. They did not report any current or previous neurological or psychiatric diseases and were free of medications. None of the participants were musicians or professional typists. Based on standardized questionnaires, participants exhibited no indication of anxiety (Beck Anxiety Inventory; (48)) or depression (Beck Depression Inventory; (49)). All participants reported normal sleep quality and quantity during the month prior to the study, as evaluated with the Pittsburgh Sleep Quality Index (50). Two participants were excluded from further analyses: one due to brain abnormality and in one participant the fMRI session was aborted prior to completion due to severe anxiety during scanning. A total of 33 participants (19 females) were considered for the analyses. Participant characteristics can be found in Supplementary Table S1. Note that data of fMRI run 8 for one participant and data of the retention test for another participant are missing.

### General experimental procedure

The experimental procedure is depicted in Figure 1. Participants were instructed to respect a regular sleep/wake schedule (according to their own schedule ± 1 hour) starting three days before the experimental session. Compliance to this schedule was assessed using sleep diaries (sleep measures are reported in Supplementary Table S1). Consumption of alcohol, nicotine, and caffeine (and other vigilance-altering substances) was prohibited the day before as well as the day of the experimental session. The St. Mary’s Hospital Sleep Questionnaire (51) was used to assess subjective sleep quality of the night before the experimental session (Supplementary Table S1). During the experimental session, participants performed different versions of an explicit serial reaction time task (SRTT; see details below and Fig. 1B; (52)). After baseline assessment of motor performance using a random SRTT, participants were trained on a sequential SRTT during which they learned a sequence of eight finger movements through repeated practice (learning session, duration: 60 min., Fig. 1A). Immediately following the learning phase, participants were placed in the MRI scanner where they performed the pre-trained sequential SRTT, as well as a random SRTT (fMRI session, duration: 60min, Fig. 1A). Last, immediately after the fMRI session, all participants completed a retention test outside the scanner (retention, duration: 10min, Fig. 1A). Both learning and retention phases took place in a behavioral testing room in the vicinity of the MRI scanner.

### Serial Reaction Time task and behavioural measures

We used an explicit bimanual serial reaction time task that was implemented in Matlab Psychophysics Toolbox version 3 (53). During the task, eight squares were presented on the screen, each corresponding to one of eight keys on the keyboard and one of eight fingers (no thumbs; Fig. 1B). The outline of the squares alternated between red and green, indicating rest and practice, respectively. During rest, participants were instructed to keep their eyes focused on the screen and their fingers still on the keyboard (duration: 10s). During practice, the outline of the squares turned green, and participants were instructed to press as quickly and accurately as possible the key corresponding to the location of a cue (green filled square) that appeared on the screen. After a response, the next cue appeared (jittered response to stimulus interval, range: 1.5-2.5s, average 2s). Depending on the task condition, the cues either followed a pseudorandom (RND; motor control task) or a fixed, repeating sequential pattern (SEQ; motor sequence learning task). In the SEQ condition, participants repeatedly practiced a pre-determined eight-element sequence in which each finger had to be pressed once (4-7-3-8-6-2-5-1, with 1 and 8 corresponding to the little fingers of the left and right hand, respectively). All participants were trained on the same sequence of movements. However, to optimize the fMRI multivariate pattern analysis (see below), the starting point of the sequence was counterbalanced across participants such that the same key was not associated to the same temporal position in a sequence across individuals (i.e., the sequence started with 4 in participant 1, with 7 in participant 2, etc.). In the RND task condition, there was no repeating sequence, but each finger had to be pressed once every eight elements. During practice of both the random and sequential SRTT, the serial position of a trial in a stream of eight elements was indicated by a green asterisk on top of the screen (Fig. 1B). These asterisks were replaced by eight red asterisks at rest.

At the beginning of the study, participants were informed that the cues could either follow a random or a repeating sequential pattern. During learning and retention, task practice took place outside the scanner, on a computer keyboard sitting at a desk, and was organized in blocks consisting of 48 keypresses each [which corresponded to six repetitions of the fixed sequence or six consecutive pseudo-random series of eight key presses], alternated with 10s rest periods (block design). During the learning session, participants completed four blocks of RND practice, followed by 20 blocks of SEQ practice. The retention test took place immediately after the fMRI session described below and consisted of four SEQ blocks followed by four RND blocks. During the fMRI session, the task was performed on an MRI compatible keyboard. Brain and behavioral data were acquired over eight consecutive scanning runs. Each fMRI run included 6 repetitions of the 8-element learned sequence (SEQ condition) and 6 pseudo-random series of 8 key presses (RND condition); thus 12 8-element key series (i.e., 6 repetitions × 2 conditions × 8 keypresses = 96 key presses/run). Repetitions of the learned sequence were randomly alternated with random patterns with the rule that a specific condition was not presented more than two times in a row. Half of the runs started with the SEQ condition, and the other half with the RND condition. In addition, three 10s rest periods were randomly inserted between 8-element series repetitions within each run. For each block (learning and retention sessions) and run (fMRI session), performance speed and accuracy were computed for each condition as the mean response time for correct responses (in seconds) and the percentage of correct responses, respectively.

### Statistical analyses of behavioral data

Statistical analyses on behavioral data were performed in SPSS Statistics 25 (IBM). To investigate behavioral changes during the learning session, for each task, a one-way RM ANOVA with block (4 for random, 20 for sequence) as within-subjects factor was performed on both speed and accuracy measures. Behavioral changes in speed and accuracy during both the fMRI and retention sessions were examined with a condition (2: sequence vs. random) by run (8) / block (4) RM ANOVA. In case of violation of the sphericity assumption, Greenhouse-Geisser corrections were applied.

### fMRI data acquisition

Images were acquired with a Phillips Achieva 3.0T MRI System and a 32-channel head coil. During the eight task runs, BOLD signal was acquired with a T2* gradient echo-planar sequence using axial slice orientation that covers the whole brain (TR =2000 ms, TE=30 ms, FA= 90°, 54 transverse slices, 2.5 mm slice thickness, 0.2 mm inter-slice gap, FoV = 210 × 210 mm^2^, matrix size = 84 × 82 × 54 slices, voxel size = 2.5 × 2.56 × 2.5 mm^3^). A structural T1-weighted 3D MP-RAGE sequence (TR = 9.5 ms, TE = 4.6 ms, TI = 858.1 ms, FA = 9°, 160 slices, FoV = 250 × 250 mm^2^, matrix size = 256 × 256 × 160, voxel size = 0.98 × 0.98 × 1.20 mm^3^) was also obtained for each participant during the same scanning session.

### fMRI data analysis

Statistical parametric mapping (SPM12; Welcome Department of Imaging Neuroscience, London, UK) was used for preprocessing of functional images and statistical analyses of BOLD data. To compute neural similarity matrices (see Representational similarity analyses section) the CoSMoMVPA toolbox was used in Matlab (54). Statistical analyses on neural similarity data were performed in SPSS Statistics 25 (IBM).

#### Preprocessing

Preprocessing included slice time correction (reference: middle slice), realignment of functional time series (to first image of each run separately in a first step and to the across-run mean functional image in a second step), co-registration of the pre-processed functional images to the structural T1-image and segmentation of the anatomical image. All the *multivariate analyses* were performed in native space (without normalization) to optimally accommodate inter-individual variability in the topography of memory representations (55). Data to be used in *univariate group analyses* (see Supplementary Methods section 1.1) were further spatially normalized to an MNI (Montreal Neurological Institute) template (resampling size of 2 × 2 × 2 mm) and spatially smoothed (functional images only, Gaussian kernel, 8 mm fullwidth at half-maximum [FWHM]).

#### Regions of interest

The goal of the fMRI analyses was to examine brain patterns elicited in specific regions of interest (ROIs) by sequence and random task practice. The following (bilateral) ROIs were selected *a priori* based on previous literature describing their critical involvement in motor sequence learning processes (e.g., (19, 24, 32)): the primary motor cortex (M1), the supplementary motor cortex (SMA), the premotor cortex (PMC), the anterior part of the superior parietal lobule (aSPL), the hippocampus, the putamen and the caudate nucleus. Cortical ROIs were defined bilaterally in MNI space and next mapped back to native space using the individual’s inverse deformation field output from the segmentation of the anatomical image. Using the Brainnetome atlas (56), M1 was defined to exclude mouth and leg representations and contained the upper limb and hand function regions of Brodmann area (BA) 4. Using the same atlas, the premotor cortex (PMC) was defined as the dorsal (A6cdl; dorsal PMC) and ventral (A6cvl; ventral PMC) part of BA 6. aSPL was defined to include anterior, medial, and ventral intraparietal sulcus of the Julich-Brain atlas (57). Lastly, the anatomical mask for the SMA was extracted from the Human Motor Area Template (HMAT; (58)). Subcortical ROIS (bilateral hippocampus, putamen and caudate) were anatomically defined in native space using FSL’s automated subcortical segmentation protocol (FIRST; FMRIB Software Library, Oxford Centre for fMRI of the Brain, Oxford, UK). All ROIs were masked to exclude voxels outside of the brain and with <10% grey matter probability and registered to the functional volumes. The average number of voxels within each ROI is reported in Supplementary Table S4.

#### Representational similarity analyses

Representational similarity analyses (RSA) were designed to investigate coding of information (1) about finger movements in their learned temporal position in the sequence (i.e., finger-position binding), (2) about temporal positions (irrespective of finger movement) and (2) about finger movements (irrespective of temporal position) within specific ROIs. As finger movements are performed in a fixed order in the sequence condition (i.e., each movement is bound to a specific temporal position), sequential SRTT data was used to identify ROIs coding for finger-position binding. As the finger-position assignment is not fixed in the random condition, random SRTT data was used to identify ROIs coding for position (irrespective of finger movement) and finger movement (irrespective of their temporal position in the pattern). For the RSA, two separate GLMs were fitted to the preprocessed functional data in each subject’s native space (see preprocessing section above). Depending on the GLM, events of interest were grouped according to temporal position or key. The first GLM (i.e., position GLM) included one regressor per temporal position (8 positions per movement pattern), separately for each condition (8 positions × 2 conditions = 16 regressors per run). The second GLM (i.e., key GLM) included one regressor for each finger/key (fingers 1 to 8), separately for each condition (8 keys × 2 conditions = 16 regressors per run). Within each regressor, events of interest (6 events per regressor per run) were modelled with delta functions (0ms duration) time locked to cue onsets. In both GLMs, key presses during rest and movement parameters (derived from realignment of the functional volumes) were modelled and entered as regressors of no interest. High pass filtering and serial correlation estimation were similar as for the univariate analyses. These GLMs generated separate maps of t-values for each regressor in each run and for each voxel of each ROI. This allowed the extraction of one across-voxel vector of t-values per regressor, run and ROI for each individual. For each voxel within each ROI, t-values were normalized by subtracting the mean t-value across all regressors (vectors) for each run separately. Note that because the order of key presses changed with every new repetition in the RND condition, an individual trial could either be classified according to its temporal position in the random pattern or according to the finger movement that was made. Accordingly, for the random condition the same events could not be modelled within the same GLM. In the SEQ condition, the same events were modelled in the position and key GLMs. As GLMs 1 and 2 included identical regressors for the SEQ condition, corresponding vectors were averaged.

Next, neural similarity matrices were estimated for each ROI. We opted for Pearson correlation as a measure of neural pattern similarity because we are primarily interested in whether the same items (i.e., positions or movements) show similarity in their neural representation, rather than in the distance between neural representations of different items. Importantly, Pearson correlation was considered a reliable measure for the purpose of this study as no differences in univariate activation were observed between conditions in our ROIs (see Supplementary Methods section 1.1 and Table S5; (59)). Neural similarity matrices were estimated by computing the Pearson correlation coefficient between vectors of t-values To do so, the full dataset (8 runs) was randomly divided into two independent subsets of 4 runs (using randperm function in Matlab). For each condition, vectors of t-values were then averaged across the 4 runs within the 2 separate subsets of runs and the resulting values were then correlated between the two datasets. This procedure was repeated 140 times (i.e., 2 conditions × 70 possible combinations when choosing 4 numbers out of 8) and correlations were then averaged across the 140 iterations, thus resulting in an 8 × 8 correlation matrix for each participant, ROI and research question of interest (see below). A total of five matrices were built to address the different research questions. As illustrated in Figure 3, three within-condition matrices were created. (1) The SEQ matrix represents pattern similarity across repetitions of the motor sequence and was used to examine brain patterns coding for the binding between finger and position in the SEQ condition. (2) The Random Position (RP) matrix represents pattern similarity across repetitions of the random patterns and was designed to examine coding for temporal position irrespective of finger movement in the RND condition. (3) The Random Key (RK) matrix was built by computing pattern similarity in the RND condition between repetitions of the same finger movement and was used to highlight coding of finger movements irrespective of their temporal position. Last, and as illustrated in Supplementary Figure S2, two “mixed’ between-condition matrices tested for similarity across SEQ and RND conditions to investigate (4) similarity of position (Mixed Position (MP) matrix) and (5) movement (Mixed Key (MK) matrix) coding between task conditions. All results of statistical analyses on the between-condition (“mixed”) matrices are reported in the supplementary material section (Supplementary Material section 2.3).

#### Statistical analyses

All correlations were Fisher-transformed before entering statistical analyses. To investigate whether a given area coded for a particular type of information, we computed for each ROI and neural similarity matrix the mean correlation in the diagonal and non-diagonal cells (27). Then, for each matrix, a (one-tailed) paired sample t-test was performed across participants to compare diagonal vs. non-diagonal mean similarity. As an exploratory follow-up analysis on the Random Key matrix, we assessed whether certain ROIs showed higher similarity between different fingers belonging to the same as compared to different hands, which would correspond to a pattern consistent with hand coding. To do so we compared for each ROI the mean correlation in the within-hand (upper left and lower right) and between-hand (lower left and upper right) quadrants of the RK matrix, while excluding cells on the primary and secondary diagonal of the matrix. To examine whether there was key beyond hand coding within the same hand, we compared the diagonal mean similarity with non-diagonal mean similarity, while excluding the between-hand quadrants. Next, to examine whether similarity changed as a function of serial position in the RND or SEQ task conditions, we ran a condition by serial position RM ANOVA on diagonal similarity values in the SEQ and RP matrices for each ROI. The results of the RM ANOVAs are reported in Supplementary Material section 2.2 and Table S2.

To assess regional coding specificity within each ROI, two separate RM ANOVAs were used to compare delta similarity (i.e., mean pattern similarity diagonal minus off-diagonal) between (1) within-condition matrices (i.e., SEQ, RP and RK; results reported in the main text) as well as between (2) between-condition matrices (i.e., SEQ, MP and MK matrices; results reported in Supplementary Material section 2.3). Lastly, for each ROI, forward stepwise linear regressions were used to identify whether finger and/or position coding (as assessed during random practice) can explain pattern similarity effects in the sequence condition (outcome: delta similarity in the SEQ matrix; predictors: delta similarity in the RK and RP matrices). Predictors were added to the model based on their contribution to the model’s explained variance and an F-test was used to assess whether the change in explained variance (i.e., R^2^ change; Δ R^2^) from the prior step was significant. If the final model obtained with the stepwise regression excluded one out of two predictors, the excluded independent variable was tested individually.

Cohen’s d effect sizes were computed for t-tests and partial eta squared (ɳ ^2^) for F-tests. For each group of hypotheses, the False Discovery Rate (FDR) procedure was applied to correct for multiple testing (i.e., two-sample t-tests corrected for 7 ROIs × 3 matrices = 21 tests for within-condition and 7 ROIs × 2 matrices = 14 tests for between-condition; ANOVA and regression results were corrected for 7 ROIs = 7 tests). In case of ANOVA, significant effects were followed up with planned pairwise comparisons and corrected for multiple comparisons using the FDR procedure. In case of violation of the sphericity assumption, Greenhouse-Geisser corrections were applied.

#### Control surrogate analyses

For each individual, surrogate neural similarity matrices were created by randomly shuffling the labels of the fingers/positions within the SEQ, RK and RP matrices (1000 permutations). Next, for each individual, the SEQ, RK and RP matrices were recalculated by subtracting their corresponding surrogate matrix (obtained by averaging over all 1000 permutations). To verify that the observed pattern similarity effects reported in the current study exceed what would be expected based on random noise (modelled in the surrogate matrices; see Fig. S5), the main analyses were repeated on these new matrices. As the pattern of results was unchanged, these results are reported in Supplementary Material section 2.4.

## Data and Code Availability

Source data for all figures, individual behavioral and pattern similarity data as well as the scripts to process the data will be made freely available on Zenodo upon publication.

### Funding

This work was supported by the Belgian Research Foundation Flanders (FWO; G099516N, 1524218N) and internal funds from KU Leuven. GA also received support from FWO (G0D7918N, G0B1419N) and Excellence of Science (EOS, 30446199, MEMODYN). ND received salary support from these grants and from the Belgian American Educational Fund (BAEF). SR is funded by a predoctoral fellowship from FWO (1141320N). HO received support from FWO (G0D3322N) and KU Leuven (C14/21/047).

## Acknowlegments

We wish to thank Ather Mujitaba for assistance with data collection and Brendan Ritchie for his advice on multi-variate analyses of fMRI data.

