## Supporting Information for "The hippocampus binds movements to their temporal position in a motor sequence"

**This PDF file includes:**

Supporting text
Figures S1 to S5
Tables S1 to S5

#### Supporting Information

### 1 Methods

#### 1.1 Univariate fMRI analyses

To assess whether there are differences in univariate activation between task conditions (SEQ SRTT vs. RND SRTT), we conducted a univariate fMRI analysis. The univariate analysis of fMRI data was conducted in two serial steps accounting for fixed and random effects, respectively. In the first-level fixed effects analysis, a general linear model (GLM) was fitted for each individual to the normalized smoothed functional data. For each run, neural responses to each cue (corresponding to a specific finger movement) in the SEQ and RND task conditions were modelled in two separate regressors using delta (stick) functions locked to the cue onset (duration 0ms, event-related design) and convolved with a canonical hemodynamic response function. Movement errors (i.e., incorrect key presses) as well as key presses during rest were modelled as events of no interest. Movement parameters (derived from realignment of the functional volumes) were entered as regressors of no interest. High-pass filtering with a cut-off period of 128 s served to remove low-frequency drifts from the time series. An autoregressive (order 1) plus white noise model and a restricted maximum likelihood (ReML) algorithm was used to estimate serial correlations in fMRI signal. Single subject contrast maps obtained from this first-level analysis and testing for the main effect of condition [RND>SEQ and SEQ>RND] were further spatially smoothed (Gaussian kernel 6 mm FWHM) and entered in a second-level random effects analysis (one sample t-test). The resulting set of voxel values constituted maps of the t statistic (SPM[T]), thresholded at p<0.05 after correction for multiple comparisons (FDR whole-brain corrected; see Table S5 for the results at the whole brain level). To verify that there were no differences in univariate activation between conditions in our ROIs, the resulting t statistic maps [SPM(T)] were masked inclusively with a mask that included all ROIs. The results are presented in Supplementary Table S5.

### 2 Results

#### 2.1 Assessment of baseline performance

Prior to learning, all participants performed a random Serial Reaction Time Task (RND SRTT) to assess baseline performance. Performance is depicted in Figure 2 of the main text. A one-way Repeated Measures (RM) ANOVA on RND SRTT performance showed that speed (reaction time) and accuracy (% correct responses) remained stable across blocks of practice (block: speed, $F_{(3,96)} = .552$, $\mu^2 = .017$, *p* = .648; accuracy, $F_{(2.228,71.286)} = .772$, $\mu^2 = .024$, *p* = .478).

#### 2.2 Pattern similarity as a function of serial position

To examine whether similarity changed as a function of serial position in the RND or SEQ task conditions, we ran a 2 (condition: SEQ vs. RND) by 8 (serial position) RM ANOVA on diagonal similarity values in the SEQ and RP matrices. Results revealed a significant position effect in all ROIs as well as a position by condition interaction in M1, PMC, SMA and aSPL (see Supplementary Table S2 for results of the ANOVAs and follow-up planned pairwise comparisons). Inspection of the data and follow-up testing indicated that position effects in the hippocampus, caudate and putamen were driven by higher similarity at the boundaries of the pattern (i.e., positions 1 and/or 8) as compared to the more central positions (2-7) irrespective of the task condition. In the SMA, aSPL and PMC, follow-up testing showed that boundary effects were more spread out in the RND as compared to the SEQ condition (i.e., in the RND condition similarity was increased for positions 1-2 as well as 7-8, while in the SEQ condition higher similarity was only observed for positions 1 and 8). Last, the interaction observed in M1 was driven by a main effect of position in the RND condition but not in the SEQ condition.

#### 2.3 Similarity in position and finger coding between conditions

Given the nature of the task, pure finger and position coding was assessed based on the random data and hence it remains unclear whether the finger and position coding as observed during random practice is similar during sequence practice. Accordingly, we designed two additional neural similarity matrices and quantified the similarity of finger and position coding between task conditions (see Fig. S2 for the structure and group average mixed neural similarity matrices). If position and/or finger coding is similar across task conditions, one would expect that the same positions or finger movements, but embedded in different task conditions, would exhibit similar activation patterns. As shown in Figure S3, analysis of the ***Mixed Key (MK) matrix*** showed that mean similarity on the diagonal (i.e., same finger) in M1 ($t_{32}$ = 16.12, d = 2.8 , $p_{FDR}$ < .001), PMC ($t_{32}$ =9.24, d = 1.61, $p_{FDR}$ < .001), SMA ($t_{32}$ = 9.14, d = 1.59, $p_{FDR}$ < .001) and aSPL ($t_{32}$= 5.35, d = .93, $p_{FDR}$ <.001) was significantly higher as compared to the off-diagonal (i.e., different finger), while no differences between diagonal and off-diagonal were found in the putamen ($t_{32}$ = .94, d = .16, $p_{FDR}$ = .176), hippocampus ($t_{32}$ = .17, d = .03, $p_{FDR}$ = .433) and caudate ($t_{32}$ =.64, d = .11, $p_{FDR}$ = .253; paired sample t-test: diagonal MK vs. off-diagonal MK). Note that no differences between diagonal and off-diagonal were expected in the caudate and the hippocampus as these ROIs showed no evidence of finger coding under random conditions. These results indicate that finger coding was similar between task conditions in the cortical ROIs.

Analysis of the ***Mixed Position (MP) matrix*** indicated that mean similarity was significantly higher for the diagonal (i.e., same position) as compared to the off-diagonal (i.e., different position) in all ROIs except the hippocampus (paired sample t-test: diagonal MP vs. off-diagonal MP; hippocampus, $t_{32}$ = -.41, d = -.07, $p_{FDR}$ = .343; caudate, $t_{32}$ = 5.95, d = .103, $p_{FDR}$ < .001; putamen, $t_{32}$ = 2.30, *d = .52,* $p_{FDR}$

< .001; aSPL, $t_{32}$= 5.56, d= .97, $p_{FDR}$ <.001; SMA, $t_{32}$ = 8.64, d= 1.5, $p_{FDR}$ < .001; PMC, $t_{32}$ = 9.78, d = 1.7, $p_{FDR}$ < .001; M1, $t_{32}$ = 8.76, d = 1.53, $p_{FDR}$ < .001; Fig. S3, grey shaded areas). To verify whether similarity effects are driven by the boundary effects reported in the main text, we removed the boundary positions and reran the analysis. The comparisons remained significant in PMC ($p_{FDR}$ = .009), caudate ($p_{FDR}$ < .001) and M1 ($p_{FDR}$ =.024), suggesting that coding of positions (other than boundaries) in these regions is shared between task conditions. It is important to note however that position coding (as assessed in the random condition) did not reach significance in M1 and caudate (see main results; paired sample t-test: diagonal RP vs. off-diagonal RP after boundary correction: M1, $t_{32}$= -1.18, d = -.205, $p_{FDR}$ = .14; caudate, $t_{32}$= .697, d = .12, $p_{FDR}$ = .27).

Next, for each ROI, delta similarity values (i.e., mean pattern similarity diagonal minus off-diagonal) were entered in a RM ANOVA with condition (SEQ, MK and MP) as within-subjects factor. The analyses revealed a significant condition effect in all ROIs (RM ANOVA; hippocampus, $F_{(2,64)}$ = 7.455, $\eta_p^2$ = .19, $p_{FDR}$ = .001; caudate, $F_{(2,64)}$ = 8.116, $\eta_p^2$ = .20, $p_{FDR}$ < .001; putamen, $F_{(2,64)}$ = 11.944, $\eta_p^2$ = .27, $p_{FDR}$ < .001; M1, $F_{(2,64)}$ = 163.917, $\eta_p^2$ = .83, $p_{FDR}$ < .001; PMC, $F_{(2,64)}$ = 104.252, $\eta_p^2$ = .76, $p_{FDR}$ < .001; SMA, $F_{(2,64)}$ = 76.664, $\eta_p^2$ = .70, $p_{FDR}$ < .001; aSPL, $F_{(2,64)}$ = 37.51, $\eta_p^2$ = .54, $p_{FDR}$ < .001). As shown in Figure S4 (panel A), planned pairwise comparisons showed that in all ROIS delta similarity in the SEQ matrix was significantly higher than in the MK (SEQ vs. MK, all $p_{FDR}$ < .001) and MP matrices (SEQ vs. MP, all $p_{FDR}$ < .01) except in the caudate where no significant difference was observed between SEQ and MP ($p_{FDR}$ = 896). These results indicate that pattern similarity is higher when information is shared within condition (as assessed in the SEQ matrix) as compared to between conditions (as assessed in the MK and MP matrices). Planned pairwise comparisons in M1 and SMA furthermore indicated that MK was significantly larger than MP (MK vs. MP, M1, $p_{FDR}$ < .001; SMA, $p_{FDR}$ = .045), while in the caudate and putamen MP was significantly larger than MK (MK vs. MP, caudate, $p_{FDR}$ < .001; putamen, $p_{FDR}$ = .044). No differences were found between MP and MK in all other regions (i.e., hippocampus, PMC and aSPL). As a follow-up, we reran the analyses after controlling for boundary effects in the SEQ and MP matrices (see Fig. S4, panel B, for corresponding results). In these analyses the main effect of condition was replicated in all regions except the caudate (RM ANOVA; hippocampus, $F_{(2,64)}$ = 5.096, $\eta_p^2$ = .14, $p_{FDR}$ = .001; caudate, $F_{(2,64)}$ = 3.018, $\eta_p^2$ = .08, $p_{FDR}$ = .056; putamen, $F_{(2,64)}$ = 9.58, $\eta_p^2$ = .23, $p_{FDR}$ < .001; M1, $F_{(2,64)}$ = 150.534, $\eta_p^2$ = .83, $p_{FDR}$ < .001; PMC, $F_{(2,64)}$ = 110.11, $\eta_p^2$ = .76, $p_{FDR}$ < .001; SMA, $F_{(2,64)}$ = 83.286, $\eta_p^2$ = .72, $p_{FDR}$ < .001; aSPL, $F_{(1.49,47.8)}$ = 36.151, $\eta_p^2$ = .53, $p_{FDR}$ < .001). As in the 8x8 matrices, pairwise comparisons showed that pattern similarity is higher when information is shared within condition (as assessed in the SEQ matrix) as compared to between conditions (as assessed in the MK and MP matrices) (SEQ vs. MP and SEQ vs. MK, all $p_{FDR}$ < .001). However, planned pairwise comparisons indicated that after controlling for boundaries MK was significantly larger than MP in all cortical regions (MK vs. MP, all $p_{FDR}$ < .001; see Fig. S4, panel B), while no differences between MK and MP were observed in the hippocampus and putamen (MP vs. MK, hippocampus, $p_{FDR}$ = .337; putamen, $p_{FDR}$ = .822). In line with the results reported in the main text, these results indicate that M1, SMA, PMC and aSPL showed preferential coding for fingers over positions, while no such differences were found in the hippocampus, caudate or putamen.

#### 2.4 Control surrogate analyses

For each individual, surrogate neural similarity matrices were created by randomly shuffling the labels of the fingers/positions within the SEQ, RK and RP matrices (1000 permutations). Next, for each individual, the SEQ, RK and RP matrices were recalculated by subtracting their corresponding surrogate matrix (obtained by averaging over all 1000 permutations). To verify that the observed pattern similarity effects reported in the current study exceed what would be expected based on random noise (modelled in the surrogate matrices; see Fig. S5), the main analyses were repeated on these new matrices and these results are reported below. In brief, these analyses confirm that all ROIs show finger-position coding in the sequence condition (beyond boundaries). They also confirm that the putamen, M1, SMA, PMC and aSPL show finger coding during random practice and that only the PMC and aSPL carry information about positions (other than boundaries) under random conditions. Also, when comparing between conditions, the control analyses essentially unchanged the results (see comparisons between representations section below).

*Finger-position binding in a motor sequence*

Analysis of the ***SEQ matrix*** revealed significantly higher mean similarity along the diagonal (i.e., same finger + position) as compared to the off-diagonal (i.e., different finger + position) in all ROIs (paired sample t-test: diagonal SEQ vs. off-diagonal SEQ; hippocampus, $t_{32}$ = 4.10, d = .71, $p_{FDR}$ < .001; caudate, $t_{32}$ = 3.87, d = .67, $p_{FDR}$ < .001; putamen, $t_{32}$ = 5.32, d = .93, $p_{FDR}$ < .001; aSPL, $t_{32}$= 10.01, d = 1.74, $p_{FDR}$ <.001; SMA, $t_{32}$ = 15.02, d =2.62, $p_{FDR}$ < .001; PMC, $t_{32}$ = 17.30, d =3.01, $p_{FDR}$ < .001; M1, $t_{32}$ = 17.52, d = 3.05, $p_{FDR}$ < .001). In line with the results reported in the main text, these results suggest that activity patterns in all ROIs carry information about the binding between finger movements and their learned temporal position in the sequence. To verify whether the binding effects are driven by boundary effects, we excluded boundary positions from the SEQ matrix and reran the analyses. In line with the results reported in the main text, these control analyses showed that pattern similarity effects in the SEQ condition (same finger + position > different finger + position) remained significant in all ROIs (paired sample t-test: diagonal SEQ vs. off-diagonal SEQ; all $p_{FDR}$ < .05).

*Finger coding in random movement patterns*

Consistent with a pattern of finger coding, analysis of the ***RK matrix*** replicated the original findings that mean similarity on the diagonal (i.e., same finger) was significantly higher as compared to the off-diagonal (i.e., different finger) in M1, PMC, SMA, aSPL and putamen (paired sample t-test: diagonal RK vs. off-diagonal RK; M1, $t_{32}$ = 18.93, d = 3.30 , $p_{FDR}$ < .001; PMC, $t_{32}$ = 10.57, d= 1.84, $p_{FDR}$ < .001; SMA, $t_{32}$ = 9.85, d = 1.71, $p_{FDR}$ < .001; aSPL, $t_{32}$= 6.46, d = 1.12, $p_{FDR}$ = .003; putamen, $t_{32}$ = 2.46, d = .43, $p_{FDR}$ < .001).

*Position coding in random movement patterns*

Analysis of the ***RP matrix*** indicated that mean similarity was significantly higher for the diagonal (i.e., same position) as compared to the off-diagonal (i.e., different position) for all ROIs with the exception of the hippocampus (paired sample t-test: diagonal RP vs. off-diagonal RP; hippocampus, $t_{32}$ = .940, d = .16, $p_{FDR}$ = .185; caudate, $t_{32}$ = 3.03, d = .53, $p_{FDR}$ = .002; putamen, $t_{32}$ = 4.33, *d = .75,* $p_{FDR}$ < .001; aSPL, $t_{32}$= 6.07, d= 1.06, $p_{FDR}$ <.001; SMA, $t_{32}$ = 5.56, d= .97, $p_{FDR}$ < .001; PMC, $t_{32}$ = 8.79, d = 1.53, $p_{FDR}$ < .001; M1, $t_{32}$ = 6.78, d = 1.18, $p_{FDR}$ < .001). To verify whether the position effects are driven by boundary effects, we excluded boundary positions from the RP matrix and reran the analyses. These control analyses showed that there is no longer evidence that activation patterns in the caudate, putamen, M1 and SMA carry information about serial positions (other than boundary positions) in the random condition (paired sample t-test: diagonal RP vs. off-diagonal RP; caudate, $t_{32}$ = .70, d = .12, $p_{FDR}$=.27; putamen, $t_{32}$ = 1.67, d = .29, $p_{FDR}$=.073; M1, $t_{32}$ = -1.18, d = -.21, $p_{FDR}$=.15; SMA, $t_{32}$ = -.81, d = -.14, $p_{FDR}$=.25). The comparison in the random condition remained significant in PMC ($t_{32}$ = 2.11, d = .37, $p_{FDR}$=.033) and aSPL ($t_{32}$ = 1.9, d = .33, $p_{FDR}$=.049).

*Comparisons between representations*

For each ROI, delta values were entered in a RM ANOVA with condition (SEQ, RK and RP) as within-subjects factor. The analyses revealed a significant condition effect in all ROIs except the caudate (hippocampus, $F_{(2,64)}$ = 5.22, $\eta_p^2$ = .14, $p_{FDR}$ = .009; caudate, $F_{(2,64)}$ = 2.73, $\eta_p^2$ = .08, $p_{FDR}$ = .072; putamen, $F_{(2,64)}$ = 6.05, $\eta_p^2$ = .16, $p_{FDR}$ = .005; M1, $F_{(2,64)}$ = 181.44, $\eta_p^2$ = .85, $p_{FDR}$ < .001; PMC, $F_{(2,64)}$ = 52.87, $\eta_p^2$ = .63, $p_{FDR}$ < .001; SMA, $F_{(2,64)}$ = 67.06, $\eta_p^2$ = .68, $p_{FDR}$ < .001; aSPL, $F_{(2,64)}$ = 17.17, $\eta_p^2$ = .35, $p_{FDR}$ < .001). Planned pairwise comparisons showed that delta similarity in the hippocampus was significantly larger in the SEQ as compared to the RK and RP conditions (SEQ vs. RK, $p_{FDR}$ = .005; SEQ vs. RP, $p_{FDR}$ = .05), while no difference was found between RP and RK ($p_{FDR}$ = .56). These data suggest that the hippocampus supports finger+position binding beyond finger or position coding. A similar pattern of results was observed in aSPL (SEQ vs. RK, $p_{FDR}$ < .001; SEQ vs. RP, $p_{FDR}$ < .001; RK vs. RP, $p_{FDR}$ = .48). In the putamen finger-position binding was greater than finger coding (SEQ vs. RK, $p_{FDR}$ = .013) but no differences were found between the other conditions (SEQ vs. RP, $p_{FDR}$ = .08; RK vs. RP, $p_{FDR}$ = .08). Planned pairwise comparisons in M1 indicated no difference in delta similarity between SEQ and RK matrices ($p_{FDR}$ = .192), but both SEQ and RK delta similarity were significantly greater than RP (both $p_{FDR}$

<.001). Lastly, in PMC and SMA a gradient in delta similarity was observed such that SEQ was greater than RK which was greater than RP (SEQ vs. RK, $p_{FDR} < .001$; SEQ vs. RP, $p_{FDR} < .001$; RK vs. RP, $p_{FDR} < .001$). In the caudate, results showed no difference in delta similarity between conditions (all $p_{FDR} > .1$). For completeness, we reran these analyses while excluding the boundary positions from the SEQ and RP matrices (as described in the previous section). The control analyses showed a significant condition effect in all ROIs except the caudate (hippocampus, $F_{(2,64)} = 4.04$, $\eta_p^2 = .11$, $p_{FDR} = .025$; caudate, $F_{(2,64)} = 1.53$, $\eta_p^2 = .05$, $p_{FDR} = .224$; putamen, $F_{(2,64)} = 4.99$, $\eta_p^2 = .14$, $p_{FDR} = .014$; M1, $F_{(2,64)} = 175.37$, $\eta_p^2 = .85$, $p_{FDR} < .001$; PMC, $F_{(2,64)} = 75.45$, $\eta_p^2 = .70$, $p_{FDR} < .001$; SMA, $F_{(2,64)} = 91.77$, $\eta_p^2 = .74$, $p_{FDR} < .001$; aSPL, $F_{(2,64)} = 15.27$, $\eta_p^2 = .32$, $p_{FDR} < .001$). The results of the planned pairwise comparisons remained unchanged as compared to the ones reported in the main text.

Altogether, these results confirm that the hippocampus and putamen show preferential coding for finger-position binding over position and finger coding. The control analyses also confirm that M1 showed preferential coding for both finger+position and finger (i.e., no difference between these two conditions) over position coding, while a gradient was observed in all the other cortical regions such that finger+position coding was greater than finger coding which was greater than position coding. Lastly, and as reported in the main text, the caudate nucleus did not show evidence of differential coding between conditions (i.e., no significant differences between conditions).

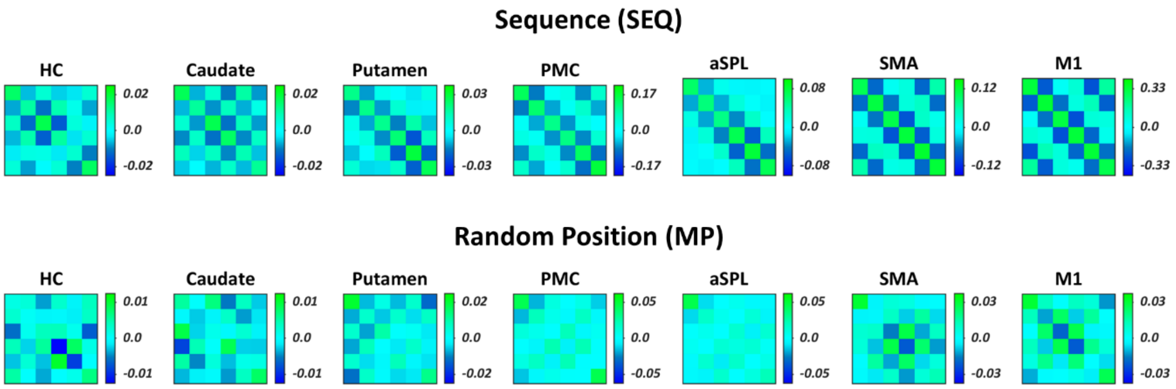


**Figure S1.** Group average within-condition neural similarity matrices for each ROI after removing the boundaries. Colour bars represent mean similarity (r). Note that colour scales are different between ROIS to accommodate for differences in signal-to-noise ratio (and therefore different effect sizes) between cortical and subcortical ROIs.

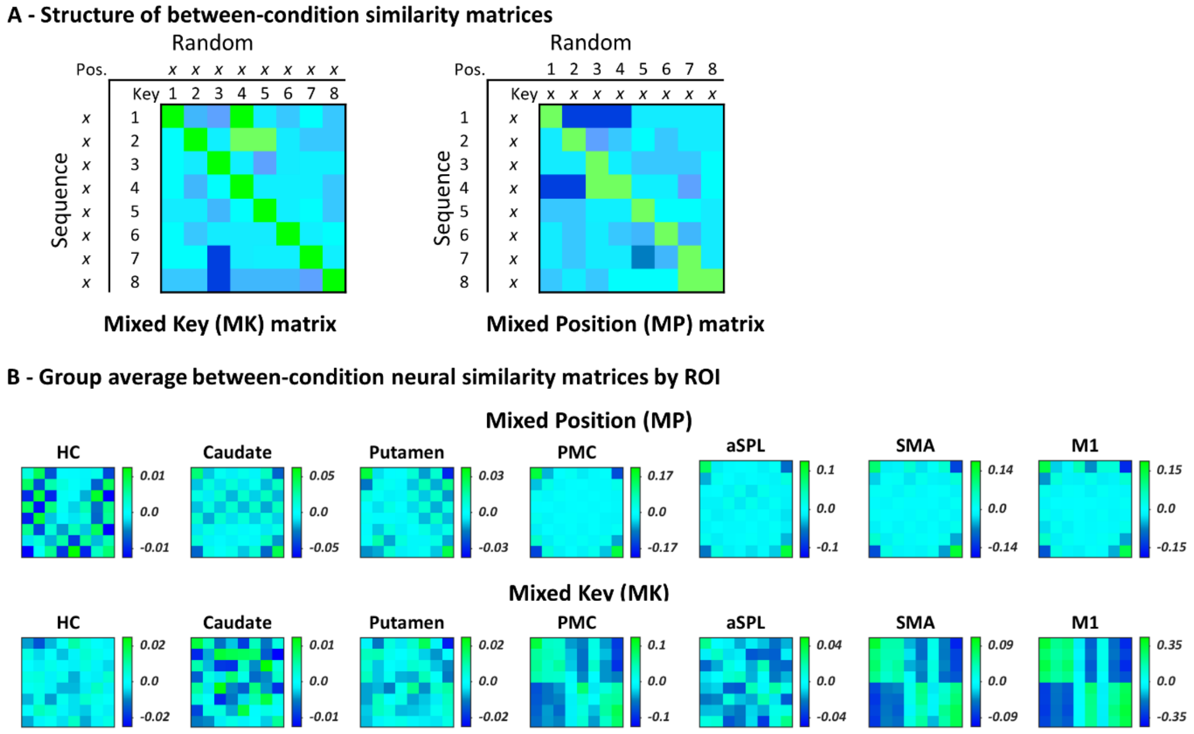


**Figure S2. (A)** Structure of between-condition "mixed" neural similarity matrices. Pattern similarity was computed across sequence and random task conditions to quantify whether finger (Mixed Key (MK) matrix) and position (Mixed Position (MP) matrix) coding was similar across conditions. In the MK matrix, diagonal cells index average similarity between the same finger movements across task conditions, while off-diagonal cells reflect average similarity across different finger movements. In the MP matrix, diagonal cells index average similarity between trials sharing the same serial position across sequence and random conditions, while off-diagonal cells reflect average similarity across different serial positions. **(B)** Group average between-condition "mixed" neural similarity matrices. Colour bars represent mean similarity (r). Note that colour scales are different between ROIS to accommodate for differences in signal-to-noise ratio (and therefore in effect sizes) between cortical and subcortical ROIs.

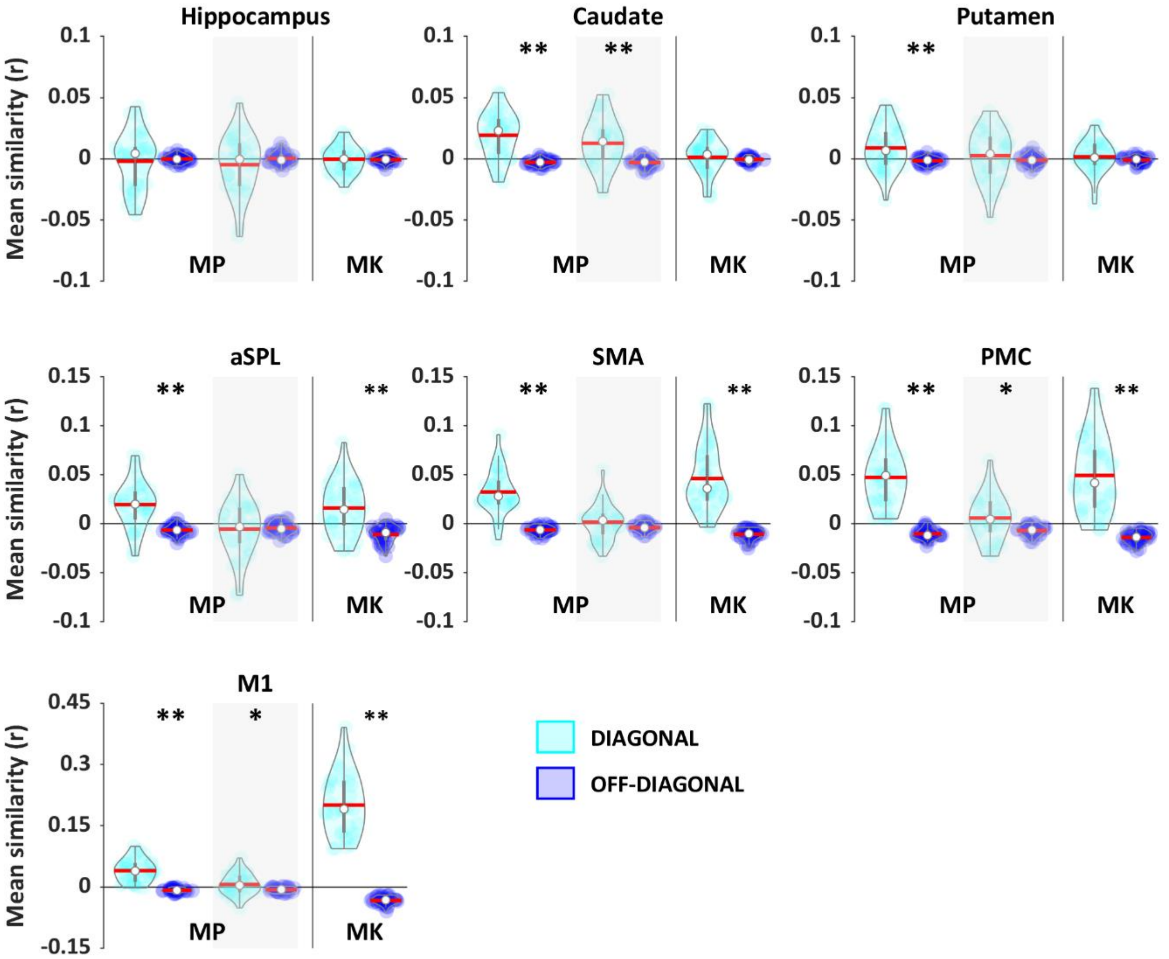


**Figure S3.** For all ROIs, mean pattern similarity for diagonal and off-diagonal cells in the between-condition "mixed" matrices (i.e., Mixed Key (MK) and Position (MP) matrices) was compared. Results show that all ROIs except the hippocampus code for position-based information similarly between sequence and random conditions. In addition, all cortical regions (aSPL, SMA, PMC and M1), but not the putamen, show similar coding of movement information across task conditions. To verify whether results in the MP matrix are driven by boundary effects, we reran the corresponding analysis without boundaries. For each ROI, results for the MP matrix after boundary correction are presented in the grey shaded areas. These analyses indicate that only the caudate, PMC and M1 show similar coding for positions across task conditions after boundary correction. It is important to note however that position coding (as assessed in the random condition) did not reach significance in M1 and caudate (see main results; paired sample t-test: diagonal RP vs. off-diagonal RP after boundary correction: M1, $t_{32}$= -1.18, d = -.205, $p_{FDR}$ = .14; caudate, $t_{32}$= .697, d = .12, $p_{FDR}$ = .27). Asterix indicate significant differences between diagonal and off-diagonal (one sided paired sample t-test; *$p_{FDR}$<.05 and ** $p_{FDR}$<.001). Coloured circles represent individual data, jittered in arbitrary distances on the x-axis to increase perceptibility. Red horizontal lines represent means and white circles represent medians. Note that Y axis scales are different between ROIS to accommodate for differences in signal-to-noise ratio (and therefore in effect sizes) between cortical and subcortical ROIs.

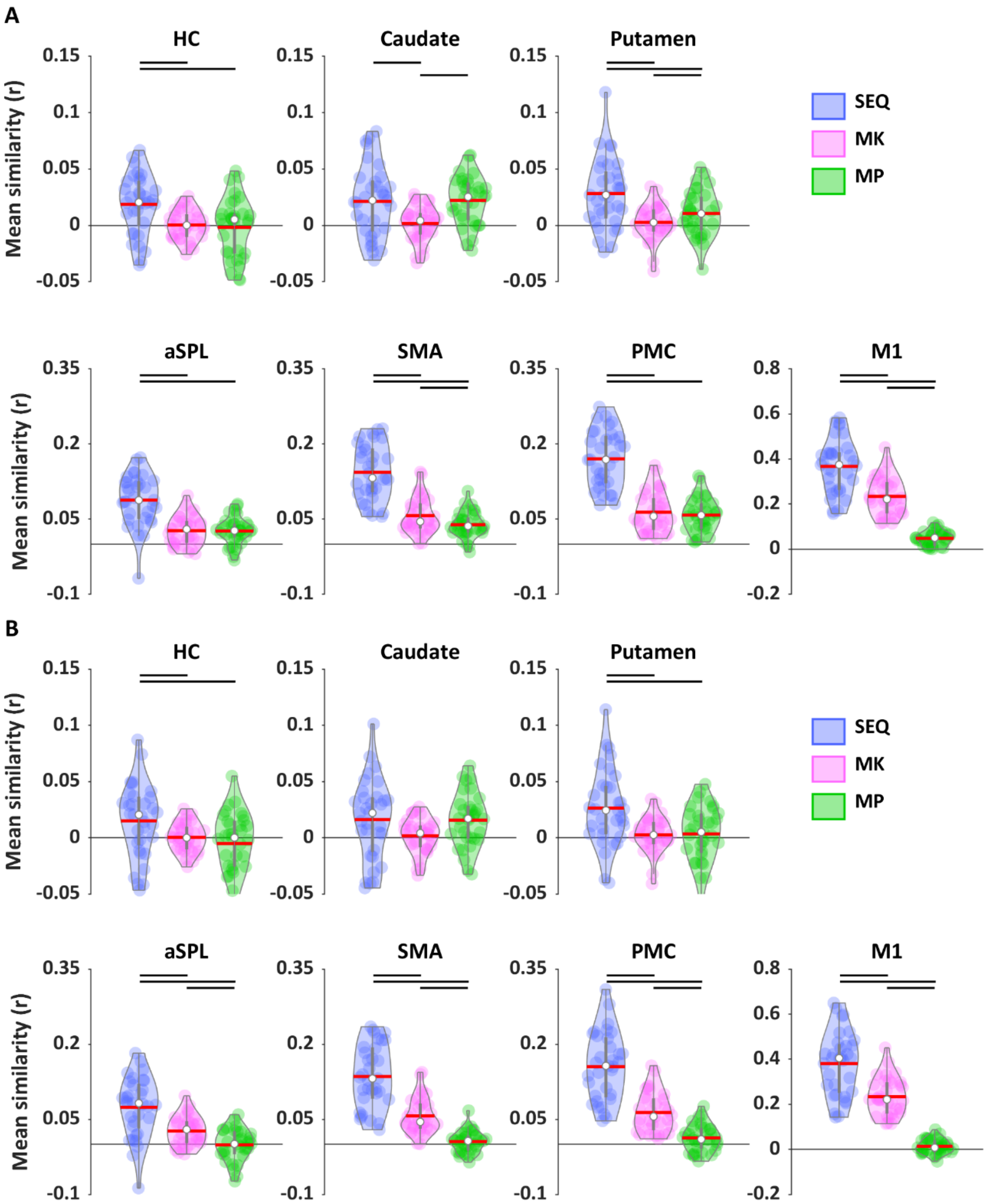


**Figure S4.** For each ROI, delta pattern similarity (diagonal minus off-diagonal) in the sequence (SEQ), Mixed Key (MK) and Position (MP) matrices **(A)** before and **(B)** after boundary correction. Pairwise comparisons showed that in all ROIs except the caudate pattern similarity is generally higher when information is shared within condition (as assessed in the SEQ matrix) as compared to between conditions (as assessed in the MK and MP matrices). Importantly, in line with the results reported in the main text, cortical regions show preferential coding for fingers over positions (but see panel A showing that in aSPL and PMC this difference is not significant before boundary correction), while in the hippocampus, caudate and putamen no such differences are found (but note that before boundary correction MP is greater than MK in striatal regions). In both panels, black horizontal lines indicate significant differences between conditions (planned pairwise comparisons following RM ANOVA) at $p_{FDR}<.05$. Coloured circles represent individual data, jittered in arbitrary distances on the x-axis to increase perceptibility. Red horizontal lines represent means and white circles represent medians.

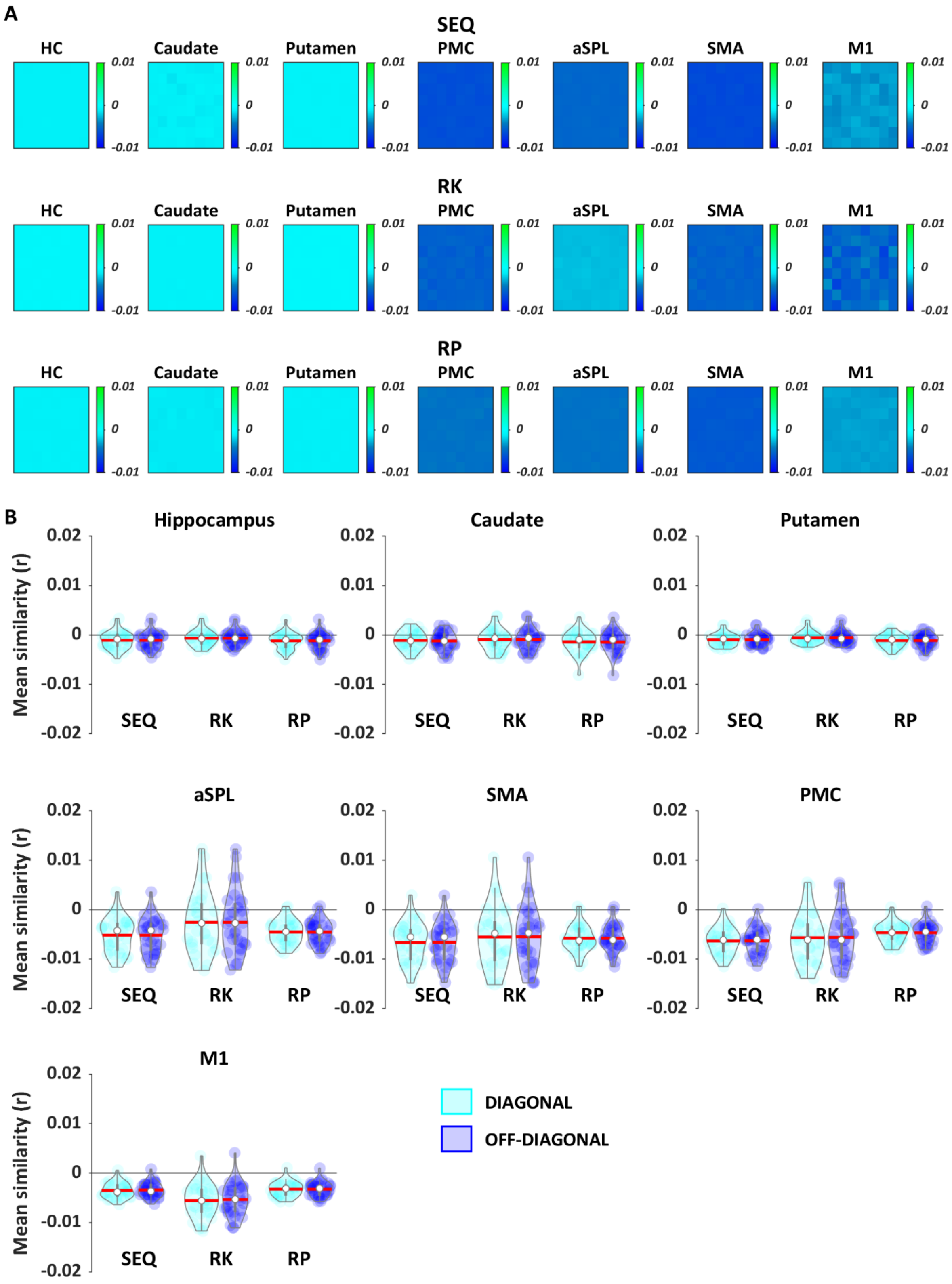


**Figure S5. (A)** Group average surrogate neural similarity matrices by condition (i.e., SEQ, RK and RP). Colour bars represent mean similarity (r). **(B)** For all ROIs, mean pattern similarity for diagonal and off-diagonal cells in the surrogate matrices by condition. There are no significant differences in mean pattern similarity between diagonal and off-diagonal cells. Coloured circles represent individual data, jittered in arbitrary distances on the x-axis to increase perceptibility. Red horizontal lines represent means and white circles represent medians.

**Table S1**. Participant characteristics

| N | 33 |
|---|---|
| Sex (F) | 20 |
| Age (yrs) | 23.4 ± 2.9 |
| Edinburgh Handedness | 0.90 ± .13 |
| Beck Depression Scale | 4.55 ± 4.78 |
| Beck Anxiety Scale | 5.39 ± 6.53 |
| PSQI[a] | 3 |
| Sleep duration Night 1 | 8hrs20min ± 53min |
| Sleep duration Night 2 | 8hrs12min ± 49min |
| Sleep duration Night 3 | 8hrs02min ± 38min |
| St. Mary's sleep quality Night 3[a] | 4 |

***Notes.*** Values are means ± standard deviation. PSQI = Pittsburgh Sleep Quality Index. [a]Median scores. Average sleep duration for each night (3 nights before the experimental session) was estimated based on the sleep diary. Subjective sleep quality of the night preceding the experimental session was assessed with the St. Mary's Hospital questionnaire.

**Table S2.** Results Condition (2: SEQ vs. RND) by Position (8) RM ANOVAs

| **Hippocampus** | **Main effect of condition** | **Main effect of position** | **Condition x Position** |
|---|---|---|---|
| | $F_{(1,32)} = 5.02$, $\eta_p^2 = .14$ $p_{FDR} = .045$ | $F_{(3.90,124.83)} = 3.66$, $\eta_p^2 = .10$, $p_{FDR} = .008$<br>*Planned pairwise comparisons:*<br>*[1 > 2-3,5-7, all $p_{FDR} < .05$]*<br>*[7 > 6, $p_{FDR} < .05$]* | $F_{(4.0\#, 128.92)} = .894$ $\eta_p^2 =.03$, $p_{FDR} = .480$ |
| **Caudate** | **Main effect of condition** | **Main effect of position** | **Condition x Position** |
| | $F_{(1,32)} = .95$, $\eta_p^2 = .03$, $p_{FDR} = .338$ | $F_{(3.27,104.83)} = 5.63$, $\eta_p^2 = .15$, $p_{FDR} = .001$<br>*Planned pairwise comparisons:*<br>*[1 >. 2-7, all $p_{FDR} < .05$]*<br>*[8 >. 2-7, all $p_{FDR} < .05$]* | $F_{(3.19, 104.83)} = 1.33$, $\eta_p^2 =.04$, $p_{FDR} = .275$ |
| **Putamen** | **Main effect of condition** | **Main effect of position** | **Condition x Position** |
| | $F_{(1,32)} = 3.38$, $\eta_p^2 = .10$, $p_{FDR} = .09$ | $F_{(3.96,127.02)} = 5.43$, $\eta_p^2 = .15$, $p_{FDR} < .001$<br>*Planned pairwise comparisons:*<br>*[1 vs. 2-6, all $p_{FDR} < .05$]*<br>*[7 > 5, $p_{FDR} < .05$]*<br>*[8 vs. 2-7, all $p_{FDR} < .05$]* | $F_{(4.36, 139.43)} = 2.0$, $\eta_p^2 =.06$, $p_{FDR} = .123$ |
| **M1** | **Main effect of condition** | **Main effect of position** | **Condition x Position** |
| | $F_{(1,32)} = 315.7$, $\eta_p^2 = .91$, $p_{FDR} < .001$ | $F_{(4.03,128.88)} = 4.28$, $\eta_p^2 = .118$, $p_{FDR} = .004$ | $F_{(4.02, 128.56)} = 7.27$, $\eta_p^2 =.19$, $p_{FDR} < .001$<br>*Planned pairwise comparisons:*<br>*RND [1 > 2-7, all $p_{FDR} < .05$]*<br>*RND [2 > 3-7, all $p_{FDR} < .05$]*<br>*RND [7 > 3-6, all $p_{FDR} < .05$]*<br>*RND [8 > 2-7, all $p_{FDR} < .05$]* |
| **PMC** | **Main effect of condition** | **Main effect of position** | **Condition x Position** |
| | $F_{(1,32)} = 80.92$, $\eta_p^2 = .72$, $p_{FDR} < .001$ | $F_{(3.6,115.03)} = 53.92$, $\eta_p^2 = .63$, $p < .001$ | $F_{(4.6, 145.91)} = 14.43$, $\eta_p^2 =.31$, $p_{FDR} < .001$<br>*Planned pairwise comparisons:*<br>*RND [1 > 2-7, all $p_{FDR} < .05$]*<br>*RND [2 > 3-6, all $p_{FDR} < .05$]*<br>*RND [7 > 3-6, all $p_{FDR} < .05$]*<br>*RND [8 > 2-7, all $p_{FDR} < .05$]*<br>*SEQ [1 > 2-5, all $p_{FDR} < .05$]*<br>*SEQ [7 > 2-6, all $p_{FDR} < .05$]*<br>*SEQ [8 > 1-7, all $p_{FDR} < .05$]* |
| **SMA** | **Main effect of condition** | **Main effect of position** | **Condition x Position** |
| | $F_{(1,32)} = 30.92$, $\eta_p^2 = .49$, $p_{FDR} < .001$ | $F_{(3.99,217.68)} = 32.26$, $\eta_p^2 = .50$, $p_{FDR} < .001$ | $F_{(4.168,133.392)} = 4.56$, $\eta_p^2 =.13$, $p_{FDR} = .004$<br>*Planned pairwise comparisons:*<br>*RND [1 > 2-7, all $p_{FDR} < .05$]*<br>*RND [2 > 3-6, all $p_{FDR} < .05$]*<br>*RND [7 > 5-6, both $p_{FDR} < .05$]*<br>*RND [8 > 2-7, all $p_{FDR} < .05$]*<br>*SEQ [1 > 4-5, both $p_{FDR} < .05$]*<br>*SEQ [8 > 1-7, all $p_{FDR} < .05$]* |
| **aSPL** | **Main effect of condition** | **Main effect of position** | **Condition x Position** |
| | $F_{(1,32)} = 12.90$, $\eta_p = 29$, $p_{FDR} = .002$ | $F_{(3.99,127.75)} = 20.04$, $\eta_p{}^2 = .39$, $p_{FDR} < .001$ | $F_{(4.59,146.77)} = 3.09$, $\eta_p^2 =.09$, $p_{FDR} = .023$<br>*Planned pairwise comparisons:*<br>*RND [1 > 2-7, all $p_{FDR} < .05$]*<br>*RND [2 > 3-6, all $p_{FDR} < .05$]*<br>*RND [8 > 2-7, all $p_{FDR} < .05$]*<br>*SEQ [1 > 3-4, both $p_{FDR} < .05$]*<br>*SEQ [8 > 1-7, all $p_{FDR} < .05$]* |

***Notes.*** In case of violation of the sphericity assumption, Greenhouse-Geisser corrections were applied. Reported effect sizes correspond to partial eta squared ($\eta_p^2$). The results of the RM ANOVAs were corrected using the False Discovery Rate (FDR) procedure for multiple testing (i.e., 7 ROIs = 7 tests). Significant effects were followed up with planned pairwise comparisons and corrected for multiple comparisons using the FDR procedure (i.e., 7 comparisons per position = 7 tests).

**Table S3.** Results linear regressions.

| | | | | | | | | | |
|---|---|---|---|---|---|---|---|---|---|
| | **Hippocampus** | **Block 1** | | | | **Block 2** | | | |
| **Stepwise** | **Predictors** | **B** | **beta** | **t** | $p_{fdr}$ | **B** | **beta** | **t** | $p_{uncorr}$ |
| | RK | .276 | 1,91 | 1.084 | .33 | .267 | .185 | 1.029 | .364 |
| | RP | | | | | .062 | .066 | .367 | 1 |
| | **Model** | $R^2$ | **F** | $\Delta R^2$ | | $R^2$ | **F** | $\Delta R^2$ | |
| | | .037 | 1.175 | .037 | | .042 | .638 | .004 | |
| | **Caudate** | **Block 1** | | | | **Block 2** | | | |
| **Stepwise** | **Predictors** | **B** | **beta** | **t** | $p_{fdr}$ | **B** | **beta** | **t** | $p_{uncorr}$ |
| | RK | | | | | -.443 | -.325 | -2.002 | .126 |
| | RP | .337 | .322 | 1.892 | .095 | .424 | .327 | .2.011 | .074 |
| | **Model** | $R^2$ | **F** | $\Delta R^2$ | | $R^2$ | **F** | $\Delta R^2$ | |
| | | .104 | 3.58 | .104 | | .209 | 3.968 | .105 | |
| | **Putamen** | **Block 1** | | | | **Block 2** | | | |
| **Stepwise** | **Predictors** | **B** | **beta** | **t** | $p_{fdr}$ | **B** | **beta** | **t** | $p_{uncorr}$ |
| | RK | | | | | -.091 | -.063 | -.352 | .848 |
| | RP | .448 | .359 | 2.144 | .07 | .474 | .380 | 2.11 | .075 |
| | **Model** | $R^2$ | **F** | $\Delta R^2$ | | $R^2$ | **F** | $\Delta R^2$ | |
| | | .129 | 4.595 | .129 | | .133 | 2.295 | .04 | |
| | **aSPL** | **Block 1** | | | | **Block 2** | | | |
| **Stepwise** | **Predictors** | **B** | **beta** | **t** | $p_{fdr}$ | **B** | **beta** | **t** | $p_{uncorr}$ |
| | RK | | | | | .045 | .027 | .149 | .882 |
| | RP | .043 | .033 | .183 | .856 | .043 | .033 | .181 | .858 |
| | **Model** | $R^2$ | **F** | $\Delta R^2$ | | $R^2$ | **F** | $\Delta R^2$ | |
| | | .001 | .034 | .001 | | .043 | .027 | .042 | |
| | **M1** | **Block 1** | | | | **Block 2** | | | |
| **Stepwise** | **Predictors** | **B** | **beta** | **t** | $p_{fdr}$ | **B** | **beta** | **t** | $p_{uncorr}$ |
| | RK | .565 | .496 | 2.637 | .021 | .517 | .455 | 2.973 | .021 |
| | RP | | | | | .976 | .266 | 1.741 | .162 |
| | **Model** | **R2** | **F** | $\Delta R^2$ | | **R2** | **F** | $\Delta R^2$ | |
| | | .246 | .003 | **.246*** | | .562 | 6.911 | .092 | |
| **Simple** | **Predictors** | **B** | **beta** | **t** | $p_{fdr}$ | | | | |
| | RP | 1.236 | .337 | 1.995 | .19 | | | | |
| | **Model** | $R^2$ | **F** | $\Delta R^2$ | | **Df 1** | **Df 2** | | |
| | | .114 | 3.979 | .114 | | 1 | 31 | | |
| | **PMC** | **Block 1** | | | | **Block 2** | | | |
| **Stepwise** | **Predictors** | **B** | **beta** | **t** | $p_{fdr}$ | **B** | **beta** | **t** | $p_{uncorr}$ |
| | RK | .647 | .573 | 3.89 | .003 | .662 | .587 | 4.156 | .007 |
| | RP | | | | | .359 | .276 | 1.954 | .21 |
| | **Model** | $R^2$ | **F** | $\Delta R^2$ | | $R^2$ | **F** | $\Delta R^2$ | |
| | | .328 | 15.126 | **.328*** | | .635 | 10.159 | .076 | |
| **Simple** | **Predictors** | **B** | **beta** | **t** | $p_{fdr\ r}$ | | | | |
| | RP | .320 | .246 | 1.413 | .168 | | | | |
| | **Model** | $R^2$ | **F** | $\Delta R^2$ | | **Df 1** | **Df 2** | | |
| | | .061 | 1.998 | .061 | | 1 | 31 | | |
| | **SMA** | **Block 1** | | | | **Block 2** | | | |
| **Stepwise** | **Predictors** | **B** | **beta** | **t** | $p_{fdr}$ | **B** | **beta** | **t** | $p_{uncorr}$ |
| | RK | .540 | .419 | 2.57 | .035 | .516 | .400 | 2.66 | .028 |
| | RP | | | | | .556 | 383 | 2.55 | .112 |
| | **Model** | $R^2$ | **F** | $\Delta R^2$ | | $R^2$ | **F** | $\Delta R^2$ | |
| | | .176 | 6.62 | **.176*** | | .323 | 7.145 | **.147** | |
| **Simple** | **Predictors** | **B** | **beta** | **t** | $p_{fdr}$ | | | | |
| | | .585 | .403 | 2.455 | .14 | | | | |
| | **Model** | $R^2$ | **F** | $\Delta R^2$ | | **Df 1** | **Df 2** | | |
| | | .163 | 6.026 | .163 | | 1 | 31 | | |

***Notes.*** *Dependent variable: delta similarity in the SEQ condition. Predictors: delta similarity in the RK and/or RP matrix. An F-test was used to assess whether the change in explained variance (*$\Delta R^2$*) from the prior step is significant (*pFDR ≤ .05, corrected for 7 tests). Block 1, df1=1, df2=31; Block 2, df1=2,df2=30. Stepwise regression was followed by simple regressions when only one out of two predictors was included in the final model.*

**Table S4.** ROI size

| **# voxels** | |
|---|---|
| M1 | Mean = 1467, *range: 1200-1722* |
| PMC | Mean = 1255, *range: 992-1486* |
| SMA | Mean = 1114,1, *range: 912-1375* |
| aSPL | Mean = 707.9, *range: 536-402* |
| Hippocampus | Mean = 852.9, range: 620 - 1028 |
| Caudate | Mean = 995.2, range: 722 – 1170 |
| Putamen | Mean = 785.7, rang: 581-952 |

*Notes.* Range across N = 33 participants.

**Table S5. Results univariate fMRI analysis**

| | | | | |
|---|---|---|---|---|
| **1. Main effect of condition [whole-brain]** | | | | |
| **[Sequence > Random] -** No suprathreshold results | | | | |
| **[Random > Sequence]** | | | | |
| R Visual association | 26 | -80 | 22 | 6.2 |
| R Angular gyrus | 44 | -62 | 18 | 5.89 |
| R Secondary Visual | 14 | -78 | -6 | 5.59 |
| **2. Main effect of condition [within ROIs]** | | | | |
| **[Sequence > Random] -** No suprathreshold results | | | | |
| **[Random > Sequence] -** No suprathreshold results | | | | |

***Notes.*** The significance threshold was set at $p_{corr} < .05$ (whole brain FDR-corrected).
